# A universal scaling law for mitotic spindles across eukaryotes driven by chromosome crowding

**DOI:** 10.1101/2025.03.05.641650

**Authors:** Lovro Gudlin, Kruno Vukušić, Maja Novak, Monika Trupinić, Matko Ljulj, Iva Dundović, Ana Petelinec, Loren Petrušić, Anna Hertel, Thomas van Ravesteyn, Marianna Trakala, Geert J. P. L. Kops, Zuzana Storchová, Josip Tambača, Nenad Pavin, Iva M. Tolić

## Abstract

Cells regulate the size of their internal structures to maintain function in diverse biological settings^1^. The mitotic spindle, a molecular micro-machine responsible for chromosome segregation^2^, must scale to accommodate genomes varying in size by over 10,000-fold across eukaryotes^3^. Yet, how spindle biomechanics adapts to vastly different genome sizes remains unknown. Here, we uncover a universal spindle scaling law, where metaphase plate width scales with genome size following a power law with an exponent of ∼1/3. We hypothesize that chromosome crowding within the metaphase plate generates compressive forces as chromosomes push against each other, thereby determining spindle size and shape. Our experiments with altered chromosome number and mechanical properties in healthy and cancerous human and mouse cells, together with a theoretical model based on inter-chromosome pushing forces and mechanical manipulations of cells, confirm this hypothesis. Extending these insights across eukaryotes, we demonstrate that chromosome crowding predicts the observed power-law scaling. The biophysical constraint of chromosome crowding offers a mechanistic explanation for the evolution of open mitosis and mitotic cell rounding, enabling the division of larger genomes. Spindle adaptability to larger genomes may promote the proliferation of polyploid cells, driving not only tumor progression but also speciation during evolution.

---

Size control is a universal regulatory principle in biology, ensuring that cellular structures scale precisely to maintain function and adaptability^1^. This challenge is particularly evident in mitosis, where the mitotic spindle must be properly scaled to divide the chromosomes accurately. Failure in this process can lead to aneuploidy, a key feature of cancer and developmental disorders^4^.

Across all eukaryotes, spindles are composed of microtubules, motor proteins, and kinetochores, which link chromosomes to spindle fibers and align them at the metaphase plate^2^. Despite their conserved molecular components, spindles have evolved diverse morphologies to adapt to different biological conditions, with pole organization varying from centrosome-focused to unfocused, kinetochore structure from monocentric to holocentric, and the mode of mitosis from closed to open^5^. In particular, spindle size is adapted to large variations in cell size during embryonic development within individual species^6^, as well as to different cell sizes and genome sizes across species^7–9^. As cells shrink during early embryonic development, spindle size decreases accordingly, regulated by mechanisms that control microtubule nucleation and dynamics in a cell size-dependent manner^10–20^.

In contrast to cell size, spindle adaptation to genome size across species has been more challenging to study due to vast and multilayered biological diversity among different organisms. Previous studies found that the genome size affects spindle geometry within a single or a few related species^21–24^, while others reported no effect^25^. Looking more broadly across all eukaryotes, it is remarkable that spindles vary in size about 100-fold, from less than 1 to 60 μm, together with the genome size variation of more than 10,000-fold^3,6,26^. Considering the extensive diversity in spindle sizes and shapes across eukaryotes, it remains uncertain whether a universal principle exists to explain how spindles handle genomes of such wide-ranging sizes.

## Spindle scaling across eukaryotes and cell ploidies

A potential evolutionarily conserved principle of spindle adaptation may be identified from the relationship between spindle and genome sizes across eukaryotes. To explore this relationship, we used published measurements or measured in published images the length and width of the spindle, as well as the width and thickness of the metaphase plate in 25 eukaryotic species from diverse lineages (Fig. 1a, b, Extended Data Fig. 1a and Table 1). For multicellular organisms with cells of varying sizes, we analyzed only differentiated cells, as they are smaller and provide less space for the same set of chromosomes than larger embryonic cells. This approach allows us to assess the lower physical limit of spindle size. Our analysis revealed that the metaphase plate width shows a sublinear relationship with the genome size (Fig. 1c). As the genome size covers 4 orders of magnitude, we plot the data on a log-log scale (Fig. 1d). Here the data align at a straight line, revealing a power-law scaling of the metaphase plate width with the genome size, with a scaling exponent of 0.33 and a significant correlation (R^2^ = 0.94, p = *10*^−*15*^). These results suggest the existence of a conserved mechanism of spindle adaptation to genome size from yeasts to animals and plants.

**Fig. 1.**
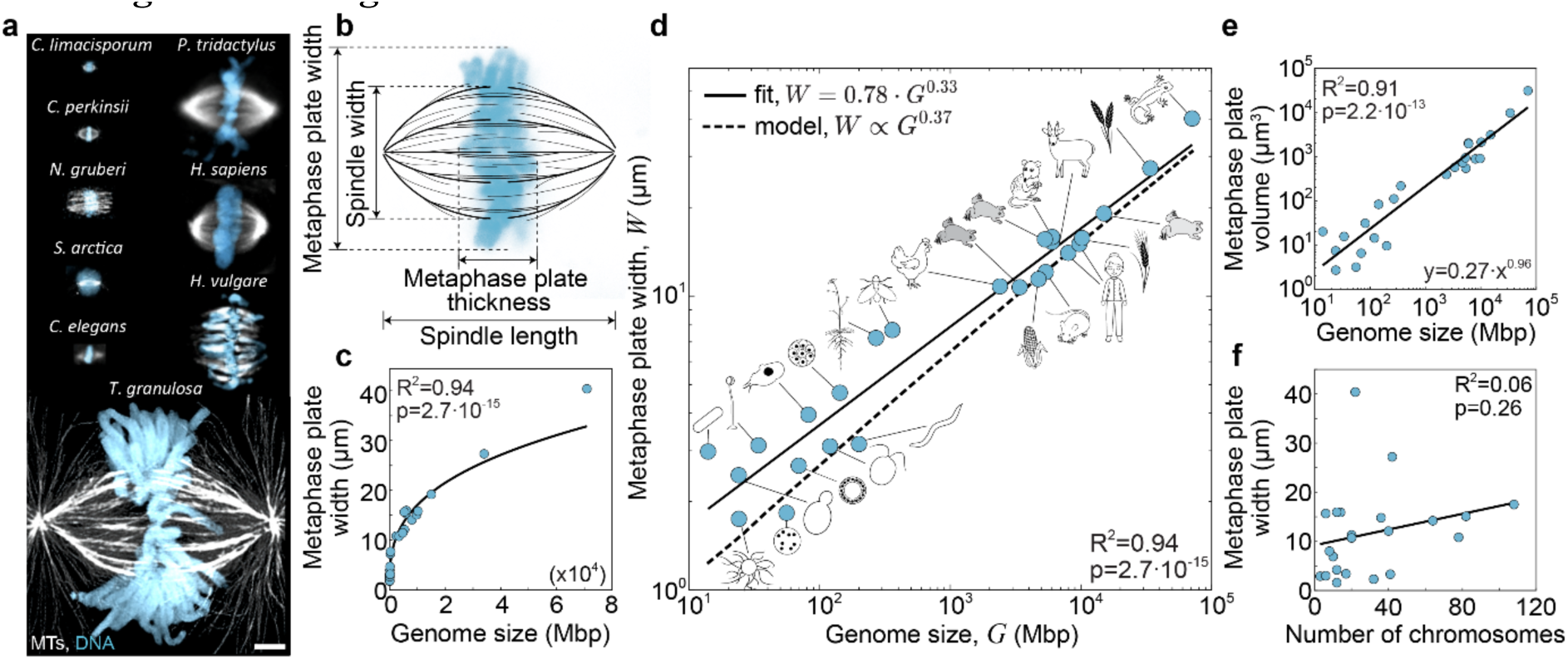
Metaphase plate width shows a power-law scaling with the genome size across eukaryotes. **a**, Spindles with labelled DNA (blue) and microtubules (gray) in denoted eukaryotic species, adapted from^21,22,117–120^. **b**, A scheme of measured spindle geometrical parameters. **c**, **d**, Metaphase plate width, *W*, as a function of genome size, *G*, for a set of eukaryotes (schematically depicted in **d**, n = 25) on a linear (**c**) and log-log scale (**d**), from **Extended Data Table 1** (note that these eukaryotes vary in the open versus closed mode of mitosis, the spindle pole structure, and the kinetochore organization, as detailed in **Extended Data Fig. 1a**). The solid line shows a power-law fit, *W* ∝ *G*^0.33^, and the dashed line is the extrapolation of the model for parameter values for human cells (Methods), where a fit yields *W* ∝ *G*^0.37^. **e**, Estimated metaphase plate volume (*V* = *πTW*^2^/4), where *T* is measured metaphase plate thickness), as a function of genome size, for a set of eukaryotes from (**d**), on a log-log scale, with a power-law fit *V* ∝ *G*^0.94^. **f**, Metaphase plate width as a function of chromosome number, *N*, for the n = 22 species from (**d**), and a linear fit. The p-values are obtained from a two-sided t-test on the regression slope. Scale bar, 5 µm.

Spindle length and width, as well as metaphase plate thickness, also show power-law scaling with genome size (Extended Data Fig. 1b-d). However, their correlation with genome size is weaker compared to that of metaphase plate width (z-score = 2.76, p = 0.0029, z-score = 2.13, p = 0.017, and z-score = 4.40, p = 5×10^-5^ for spindle length, width, and metaphase plate thickness, respectively). Together, these relationships identify metaphase plate width as the key spindle parameter scaling with genome size.

Previous analyses of condensed metaphase chromosomes revealed a remarkably uniform chromatin density across human chromosomes and a subset of eukaryotic species (∼90 Mbp μm^-3^)^27–29^. To assess whether this constancy extends to the scale of the entire metaphase plate, we estimated plate volume from measured width and thickness across diverse eukaryotes (Methods). Plate volume scaled with genome size according to a power law with an exponent of 0.96 (R² = 0.91, p = 10^-13^; Fig. 1e), indicating a near-linear increase in plate volume with chromatin amount. Accordingly, effective metaphase plate density (6.27±0.99 Mbp μm⁻³) was roughly constant across genome sizes (Extended Data Fig. 1e). The lower effective density of the whole metaphase plates relative to isolated chromosomes^27–29^ likely reflects inclusion of inter-chromosomal spaces in plate volume estimates. Plate shape was likewise similar across eukaryotes, with a width to thickness aspect ratio of 1.89 ± 0.59 (Extended Data Fig. 1f, g). Together, these results indicate that metaphase plate density and shape are conserved across species.

To explore the role of chromosome number in the observed scaling, we examined the scaling behavior both across varying chromosome numbers and when chromosome number was held approximately constant. Across all species, metaphase plate and spindle dimensions showed no correlation with chromosome number, in contrast to their strong correlation with genome size (Fig. 1f, and Extended Data Fig. 1h-j). Focusing on a subset of species with similar chromosome numbers, ranging from 32 in yeast to 42 in wheat (Extended Data Table 1), we found that metaphase plate width remained strongly correlated with genome size despite the narrow range of chromosome numbers (R² = 0.97, p = 0.0028). We infer that the total amount of chromatin in the metaphase plate, rather than chromosome number, underlies the observed scaling.

To uncover the physics behind the identified scaling law, we start by exploring the interplay between the mechanics of the mitotic spindle and chromosomes, which are aligned at the metaphase plate by microtubules and motor proteins^30,31^. We hypothesize that crowding of chromosomes, as they “fight for space” within the metaphase plate, results in inter-chromosome pushing forces which increase the plate and spindle width. This hypothesis is supported by the tight packing of chromosomes within the metaphase plate^32^, where they repulse each other by steric forces arising from chromosome compaction and by electrostatic interactions^33–36^. Alternatively, spindle geometry may be set by self-organization of the microtubule network^25,37^. To test the chromosome-crowding hypothesis, we used four complementary approaches that alter chromatin amount and mechanical properties, and perturbed microtubules to evaluate the alternative hypothesis.

First, we increased cell ploidy to raise the total amount of chromatin by increasing chromosome number. We used hypotetraploid clones derived from human non-transformed RPE1 and tumor HCT116 cell lines, which have undergone long-term adaptation after cytokinesis failure and divide in a bipolar manner^38,39^ (Fig. 2a, and Extended Data Fig. 2a-c). To generate newly-formed tetraploid and hypooctaploid cells with bipolar spindles, we designed novel strategies by preventing centriole overduplication^40^ after induced polyploidization through mitotic slippage^41^ or cytokinesis failure^42^ (Extended Data Fig. 2a, d).

**Fig. 2.**
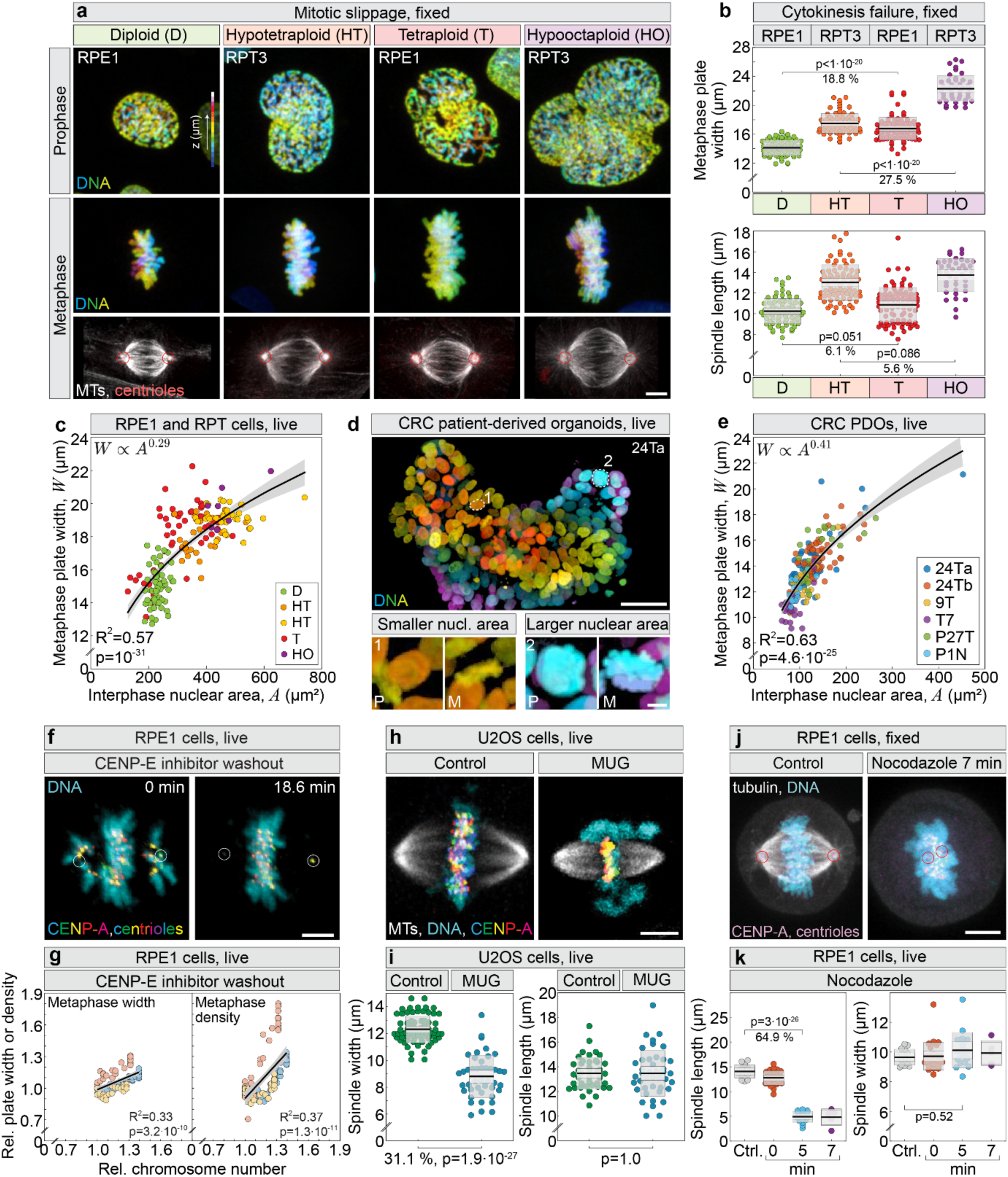
Sublinear scaling of metaphase plate width with ploidy in differentiated cells and patient-derived organoids. **a**, DNA in RPE1 and RPT3 cells of different ploidy after polyploidization by mitotic slippage expressing H2B-GFP during prophase (top), and metaphase (middle), with immunostaining for α-tubulin (MTs, gray), and centrin-3 (centrioles, red, encircled) (bottom). **b**, Metaphase plate width (top) and spindle length (bottom) measured in cells of different ploidy after polyploidization by cytokinesis failure. Number of cells: 86, 76, 94, 42. Colored points represent individual cells; black lines show the mean, with light and dark gray areas marking 95% confidence intervals for the mean and standard deviation, respectively. Percentages indicate differences between groups. **c**, **e**, Metaphase plate width, *W*, versus interphase nuclear area, *A*, in cells of different ploidies (**c**), and in colorectal cancer (CRC) patient-derived organoids (PDOs) lines (**e**), each obtained from independent patient tumor samples (24Ta, 24Tb, 9T, T7, P27T, where “T” indicates tumor) and control sample (P1N, where “N” indicates normal)^47^, with a power-law fit and 95% confidence intervals. In (**c**), orange dots represent RPT3 and yellow RPT1 cells. Number of cells: 58, 28, 36, 51, 7 (c); 58, 59, 21, 27, 23 (e). **d,** CRC PDOs expressing H2B-GFP (top), with cells of smaller (1) and larger (2) ploidy enlarged (bottom). Movies obtained from^47^. **f**, Live-imaged diploid RPE1 cells expressing CENP-A-GFP and centrin-3-GFP (color coded for depth with circled centrioles), and stained with SPY650-DNA (cyan), after washout of the CENP-E inhibitor. Time after the washout is given. **g**, Relative metaphase plate width (left) and relative chromatin density (right) in all analyzed time frames versus relative number of chromosomes, after acute reactivation of CENP-E. Black line, linear fit; gray area, 95% confidence interval. Number of cells: 3 (color coded dots) from three independent experiments. Values were calculated relative to the initial value at the start of imaging. **h**, **i**, U2OS cells expressing CENP-A and α-tubulin (MTs) during control mitosis (top) and mitosis with an unreplicated genome (MUG, bottom), with spindle width (top) and length (bottom) quantified for both conditions (**i**). Number of cells: 12, 11 (top); 12, 11 (bottom). DNA and CENP-A are color-coded for depth. **j**, Control (left) and nocodazole-treated (7 min, right) metaphase RPE1 cells expressing CENP-A-GFP and centrin-GFP (magenta), fixed and immunostained for α-tubulin (MTs, gray), with DNA labelled with DAPI (cyan). **k**, Spindle length (left) and spindle width (right) for control and nocodazole-treated cells at different times after the treatment. Number of cells: 16 (control) and 29 (nocodazole). Abbreviations: Ctrl., control; Rel., relative; nucl., nuclear. Scale bars, 5 µm. **b**, **i,** and **k**, data are mean with 95% confidence intervals and standard deviation. **b**, **k** Tukey’s Honest Significant Difference test. **c**, **e**, **g**, **i**, Student’s two-sided t-test. Each with data was obtained from at least three independent experiments.

Total chromosome signal intensity increased nearly twofold upon ploidy doubling (Extended Data Fig. 2e), confirming successful polyploidization. We found that metaphase plate width increased with ploidy across all cell types and polyploidy-induction methods, whereas metaphase plate thickness, spindle length, and spindle width increased in most, but not all, cases (Fig. 2b, Extended Data Fig. 2f-i, and Movie 1). The spindle or metaphase plate widening following polyploidization was not dependent on the presence of centrioles at the spindle poles in RPE1 cells (Extended Data Fig. 3a-c), as well as in naturally acentriolar mouse embryonic cells^43^ (Extended Data Fig. 3d, Table 2). Together, these results demonstrate that the metaphase plate width scales robustly with chromatin content upon polyploidization.

In polyploid cells, we reasoned that the higher chromosome number would increase chromatin density within the metaphase plate, reflecting chromosome crowding and the resulting inter-chromosome pushing forces. Indeed, chromatin density increased after ploidy doubling by 19% in tetraploid and 17% in hypooctaploid RPE1 cells (Extended Data Fig. 3e). In parallel, metaphase plate width and thickness increased by 19% and 28% in tetraploids, and by 29% and 37% in hypooctaploids (Fig. 2b, Extended data Fig. 2f). The overall increase over all ploidies was higher for the plate dimensions than for chromatin density. These data indicate that while higher ploidy modestly elevates chromatin density, cells primarily accommodate additional chromatin through expansion of the metaphase plate volume.

To test the relationship between the metaphase plate width and ploidy on a cell-by-cell basis, we used the nuclear area in prophase as an established proxy for the ploidy in each cell^41^. The metaphase plate width in live-imaged cells correlated with the nuclear area and a power-law fit yielded an exponent of 0.29 (Fig. 2c), matching the scaling found across eukaryotes (Fig. 1d). In fixed cells, metaphase cell area was used as an indirect proxy for ploidy because it increased markedly with ploidy (Extended Data Fig. 3f). Consistent with this, metaphase plate width showed a similar power-law scaling with cell area as with nuclear area (Fig. 2c, Extended Data Fig. 3g), indicating that the scaling relationship is robust to the choice of ploidy proxy.

Increased ploidy affects not only chromosome number and cell size, but also the transcriptome^44,45^. Indeed, our transcriptome analysis showed upregulation of key spindle assembly and kinesin family genes (e.g., AURKB, AURKA, KIF4A) in adapted tetraploid cells (Extended Data Fig. 4a-c), but to a lesser extent than the twofold increase corresponding to doubling of ploidy, indicating that mitotic factors increase sublinearly with ploidy^46^.

In the second approach, we took advantage of natural variation in ploidy in patient-derived colorectal cancer organoids^47^ (Fig. 2d). Metaphase plate width displayed a strong correlation with nuclear area, exhibiting a power-law scaling (Fig. 2e) consistent with that observed in the RPE1 cell line and its ploidy-diverse derivatives (Fig. 2c). In contrast, metaphase plate thickness showed a weak correlation with nuclear area across both systems (Extended Data Fig. 3h, i). In agreement with these results, naturally occurring polyploids among primary mouse small intestinal cells grown in 3D^48^ had wider metaphase plates, as well as wider and longer spindles (Extended Data Fig. 3j, k).

Third, we reduced the number of chromosomes in the metaphase plate by perturbing CENP-E/kinesin-7 in otherwise diploid RPE1 cells, resulting in hypodiploid metaphase plates with a few chromosomes at the spindle poles, which could be precisely quantified in individual cells (Fig. 2f, Extended Data Fig. 5a). Based on the chromosome crowding hypothesis, we predicted that metaphase plate width and chromatin density would increase as additional chromosomes entered the plate following acute reactivation of CENP-E^49^. Indeed, as chromosomes progressively accumulated within the metaphase plate, both plate width and chromatin density increased, accompanied by an increase in spindle width and a decrease in plate thickness (Fig. 2g, Extended Data Fig. 5a-c, Extended Data Movie 3). Similar trends were observed in fixed CENP-E-depleted cells (Extended Data Fig. 5d, e) and in published images of near-haploid human tumor HAP1 cells^50^, which displayed metaphase plates that were 18% narrower than those of near-diploid cells (Extended Data Table 2). Together, these observations demonstrate that inter-chromosome pushing forces, reflected by changes in metaphase plate width and chromatin density, increase as the number of chromosomes within the plate rises.

Fourth, we reduced the amount of chromatin in the metaphase plate and disrupted chromosome structure by inducing mitosis with an unreplicated genome (MUG)^51^ in the U2OS osteosarcoma cell line (Fig. 2h). In MUG cells, where centromeres are aligned on the spindle but most of the uncondensed chromatin is displaced in the cytoplasm, spindles were 31% thinner without changes in their length or cell size (Fig. 2i; Extended data Fig. 5d, e; note that the metaphase plate width cannot be determined due to uncondensed chromatin). These experiments indicate that the amount of chromatin and its mechanical properties determine spindle width.

To investigate whether the observed spindle scaling with ploidy in mitotic cells is conserved in meiotic cells, we analyzed published images of mammalian meiotic spindles. Consistent with the chromosome crowding hypothesis, metaphase I plates, having twice the ploidy of metaphase II plates, were 23% wider in human oocytes and 33% wider in mouse oocytes in comparison with metaphase II plates (Extended Data Table 2)^52–57^. Despite the variation in plate width, spindle length differed by no more than 7% between metaphase I and II (Extended Data Table 2). Similarly, tetraploidization increased the width of the metaphase plate by 26% in porcine oocytes^58^, while having a significantly smaller impact on spindle length (Extended Data Table 2). Collectively, our experiments and analyses support the hypothesis that total chromatin amount, and the resulting degree of chromosome crowding within the metaphase plate, sets spindle and metaphase plate width in mitosis and meiosis across diverse culturing conditions, cell types, and species.

To test the alternative hypothesis that spindle width is maintained by microtubule-generated forces^25,37^, we acutely depolymerized microtubules in the metaphase spindle in diploid RPE1 cells using 1 μM nocodazole (Fig. 2j, Extended Data Fig. 6a, Extended Data Movie 4). Nocodazole caused a rapid and pronounced reduction in spindle length (65%), whereas spindle width and metaphase plate width remained unchanged over 7 minutes, and control metaphase cells showed no changes (Fig. 2k, Extended Data Fig. 6b, c). Immunofluorescence confirmed near-complete microtubule depolymerization within ∼7 minutes of nocodazole treatment (Extended Data Fig. 6e, f). These observations show that metaphase spindle width is maintained primarily by the physical properties of metaphase chromatin and inter-chromosome pushing forces.

## Inter-chromosome pushing forces set spindle size

To directly test the chromosome crowding hypothesis and estimate the resulting inter-chromosome pushing forces, with the ultimate aim of explaining the spindle scaling law, we introduced a physical model of the spindle. We chose a simple approach where the spindle is composed of chromosomes and microtubules, while omitting proteins unrelated to the chromosome crowding hypothesis, such as microtubule-associated proteins (MAPs). We model each chromosome as a sphere because this is the simplest geometric object characterized solely by its radius, and we use the sphere volume as a proxy for the amount of chromatin (Fig. 3a, and Box 1). In our model, the chromosomes are compressible in response to forces, based on previous measurements of chromosome elasticity^59^. They are attached to microtubules and positioned in the spindle midplane, representing the metaphase plate. The microtubules, which extend between the spindle poles, push the chromosomes inward towards the spindle axis through their bending forces (Fig. 3a). In response, chromosomes exert inter-chromosome pushing forces as they squeeze against one another. Together, this balance of microtubule bending forces and inter-chromosome pushing forces defines the mechanical equilibrium that sets the chromosome positions and thereby the width of the metaphase plate and the overall spindle architecture (Box 1, Methods).

**Fig. 3.**
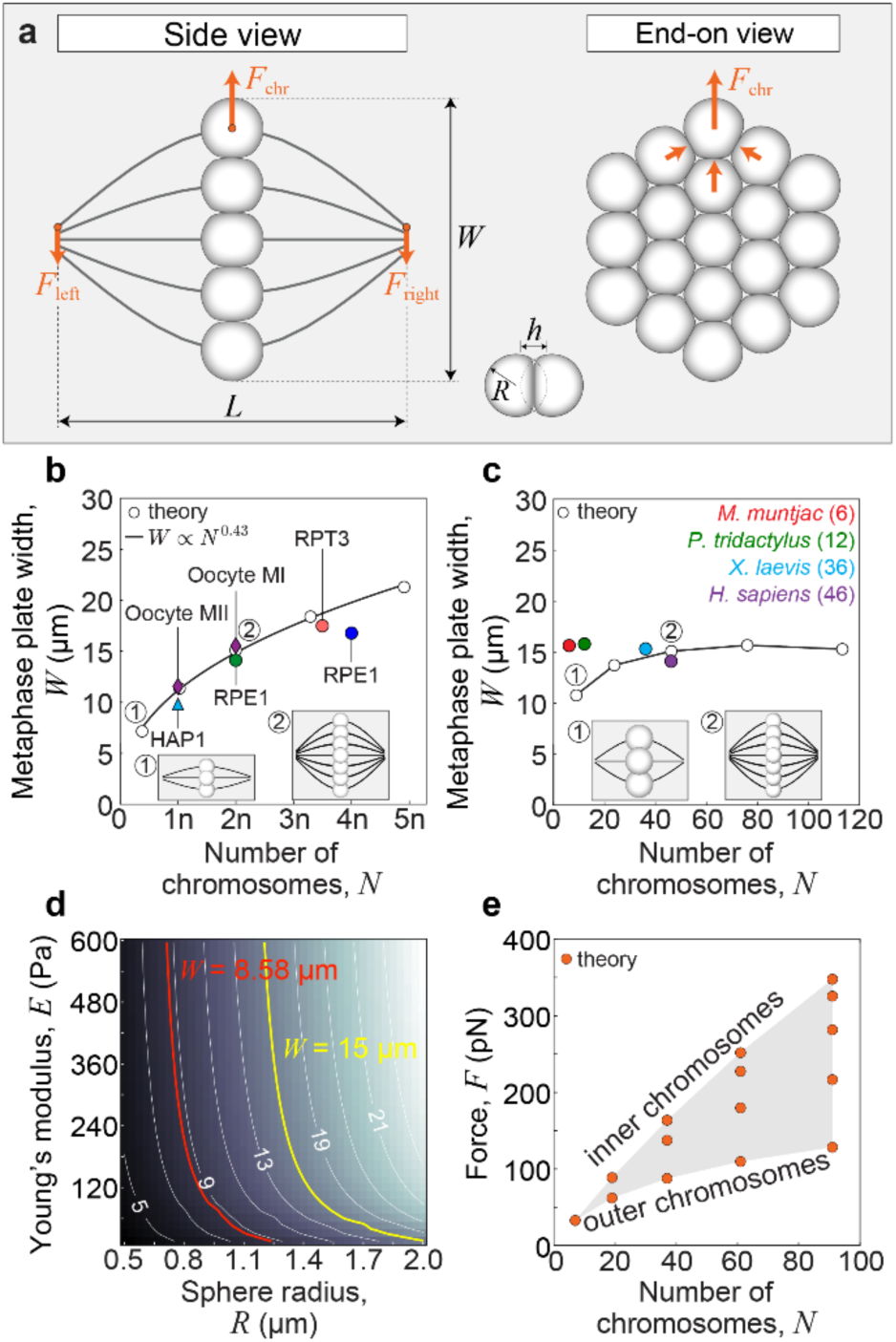
Physical model of the metaphase plate. **a**, Geometry of the metaphase spindle (spindle length, *L* and metaphase plate width, *W*) with 19 chromosomes (deformed spheres) and forces exerted on the individual microtubule bundle (orange arrows), *F*_left_, *F*_right_, and *F*_chr_, shown in the side view (left) and end-on view (right). Force at the chromosome, *F*_chr_, arises from the repulsion of its nearest neighbours. Positions of spheres are calculated by the model, whereas their deformations are schematically represented, see example of two spheres in contact with labelled reduction in distance between the sphere centres, ℎ, and the radius of each sphere, *R*. **b**, Metaphase plate width versus the chromosome number and the fit. The value of lattice parameter is *α*_0_ = 0.0025. Colored dots represent experimentally measured means of metaphase plate widths for: near-haploid HAP1 cells (Extended Data Table 2), diploid metaphase I (MI) and haploid metaphase II (MII) human oocytes (Extended Data Table 2), and diploid and tetraploid RPE1, as well as hypotetraploid RPT3 cells (Fig. 2, Extended Data Fig. 2). **c**, Metaphase plate width versus the chromosome number and the results of the model. The radius of the sphere and chromosome number are varied simultaneously, whereas the total spindle volume is held constant and consistent with parameter values *N* = 37 and *R* = 1.25 μm. Colored points represent experimentally measured means of metaphase plate widths for given species, whose genome size is 5–8 Gbp (Extended Data Table 1). Insets (1) and (2) show schematic spindles with 7 and 37 chromosomes, respectively (**b** and **c**). **d**, Phase diagram of metaphase plate width as a function of the chromosome’s Young’s modulus and chromosome radius. The yellow and red lines show values of *R* and *E* yielding *W* = 15 μm and *W* = 8.58 μm, respectively, whereas the white lines correspond to the spindle width values shown in the plot. **e**, Inter-chromosome pushing force versus the chromosome number calculated as the total force exerted by 3 inner neighbors, as depicted in (**a**). If not stated otherwise, parameter values are: Young’s modulus *E* = 400 Pa, Poisson ratio *σ* = 0.3, *N* = 37, *L* = 14 μm, *R* = 1.25 μm, and *α*_0_ = 0.25 μm.

Motivated by our experiments on altered ploidy and chromosome structure (Fig. 2), we use our model to explore how chromosome number, size, and stiffness affect the inter-chromosome pushing forces and the resulting spindle shape. In an initial step to describe a baseline spindle, our model reproduces a typical spindle shape and predicts only slight deformation of chromosomes (Fig. 3a), for the known stiffness of chromatin and microtubules^60,61^. We next alter the chromosome number while keeping constant either the volume of individual chromosomes or their total volume. In the case of constant volume of individual chromosomes, where an increased number of chromosomes corresponds to the increased total amount of chromatin, the model predicts a power-law scaling of the metaphase plate with an exponent of 0.43 (Fig. 3b). In contrast, in the case of constant total volume of chromosomes, where individual chromosomes become smaller as their number increases, the model predicts a roughly constant metaphase plate width except for narrower plates at small chromosome numbers (Fig. 3c). Thus, spindle shape is set primarily by the total chromosome volume rather than by their number.

To test the model’s predictions, we directly compare them with the experimental results on different ploidy. Measurements of spindles in cells with altered ploidy from haploid to diploid to tetraploid (Fig. 2, Extended Data Fig. 2 and 5, Extended Data Table 2), and in oocytes that naturally change from diploid to haploid state (Extended Data Table 2), follow the model’s prediction that doubling of chromosome number results in a 35% wider metaphase plate (Fig. 3b). To test the case of constant total volume of chromosomes, we compared 4 species that have a similar genome size (5–8 Gbp), but a substantially different chromosome number (6– 46, Extended Data Table 1). These species have a similar metaphase plate width, in agreement with a roughly constant width predicted by the model (Fig. 3c). Thus, irrespective of the way the chromosome number was altered experimentally, and even when comparing different species, the experimental data agree with the model’s prediction that the spindle shape is determined by the total chromosome volume.

To explain the experiments on MUG cells, which have largely uncondensed chromosomes and narrower spindles (Fig. 2h, i), we use the model to explore how spindle width depends on chromosome stiffness together with size. The model predicts narrower spindles for softer or smaller chromosomes (Fig. 3d). This theoretical result suggests that MUG experiments can be interpreted by alterations of chromosome mechanical properties and size.

Beyond interpreting the experiments, the model gives predictions on inter-chromosome pushing forces, which are difficult to access experimentally. We find that these forces are in the range of hundreds of pN (Fig. 3e), comparable to the pulling forces acting on kinetochores^62–64^. As the number of chromosomes increases, the inter-chromosome pushing forces also increase. The outermost chromosomes experience the lowest net force because they have fewer neighboring chromosomes pushing on them. In contrast, as we move toward the center, each chromosome is subjected to progressively stronger forces, resulting from the increasing number of chromosomes collectively pushing inward and amplifying the overall compressive load (Fig. 3a, e).

To test under what conditions the inter-chromosome pushing forces are large enough to displace the chromosomes from the metaphase plate, we performed stability analysis of the metaphase plate in the extended model where chromosomes are aligned at the spindle midplane by an elastic spring (Methods). We obtain that chromosome alignment at the metaphase plate is stable if the spring is stiff enough, which is the case for diploid human cells (Extended Data Fig. 2f-i). In contrast, in cells with higher ploidies, the metaphase plate can become unstable for the same spring stiffness, in agreement with the experimentally increased thickness of the metaphase plate (Extended Data Fig. 2f-i).

As our central hypothesis relies on chromosome and microtubule mechanics, its key test involves acutely perturbing mechanical forces in the spindle, such as through external compression, rather than altering spindle proteins, given their multiple roles in mitosis^65^. The main prediction of the model is that applying compressive force from the top causes an increase in inter-chromosome pushing forces and widening of the metaphase plate, accompanied by a reduction of height and changes of microtubule angles due to the freely jointed ends (Fig. 4a-d, and Extended Data Fig. 7a, b). A significant deformation of the metaphase plate occurs as the spheres redistribute under force, with those in the central region undergoing substantial deformation (Fig. 4a). Spindles with smaller or softer spheres also become deformed under compression, but to a smaller extent (Fig. 4c, Extended Data Fig. 7a, b).

**Fig. 4.**
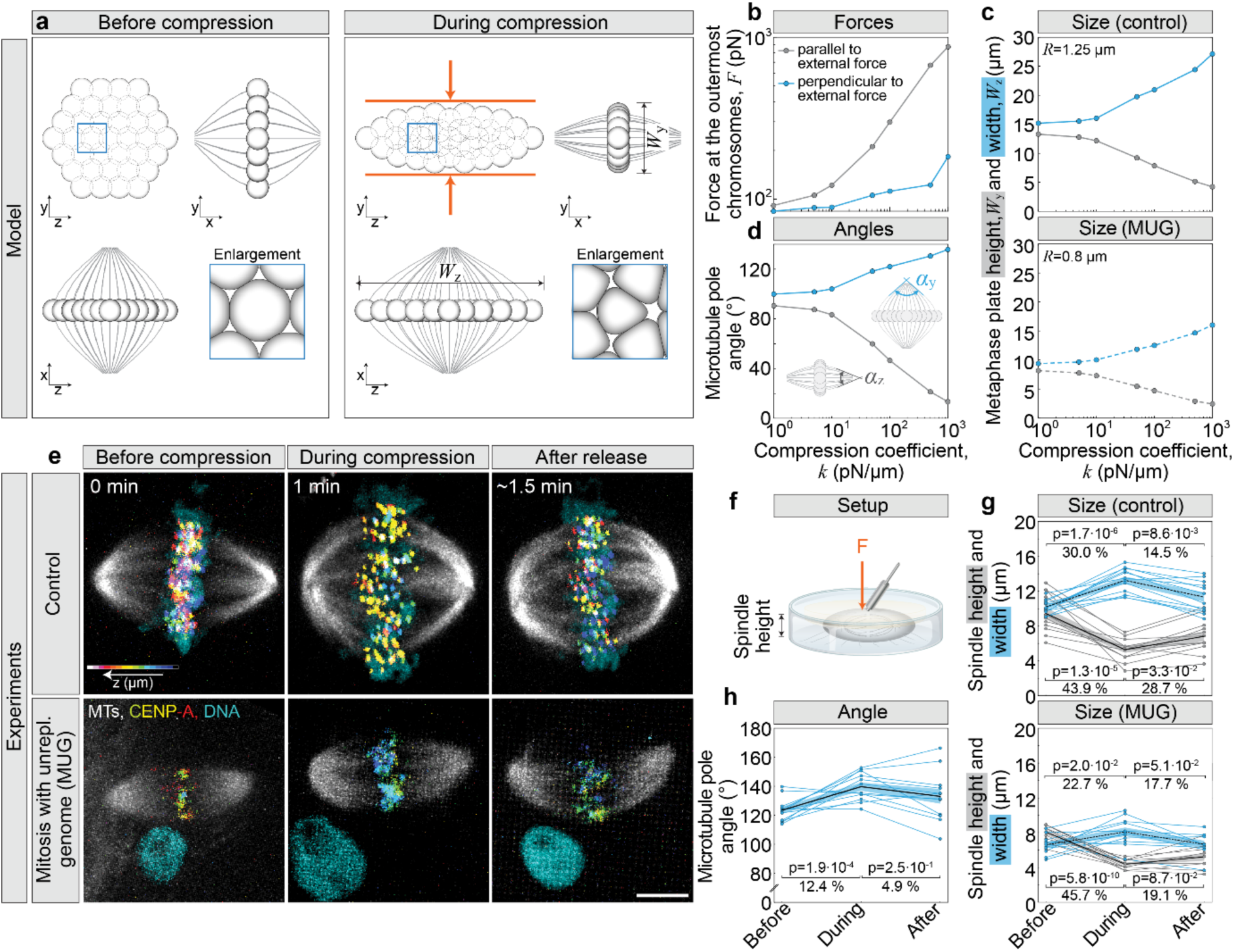
Inter-chromosome pushing forces within the metaphase plate cause spindle expansion. **a**, Examples of two solutions of the model without (left) and with (right) the external force exerted at the chromosomes, depicted by red arrows. Compression coefficient is *k* = 100 pN/µm. Solutions are shown in the end-on view (upper left), and two side views (upper right, metaphase plate height, *W*_*y*_, and lower left, metaphase plate width, *W*_*z*_). **b**, Inter-chromosome pushing force at the outermost chromosomes, at *y* = 0 and *z* = 0, versus compression coefficient, *k*. **c**, Model prediction for the metaphase plate width and height versus compression coefficient, for two different sphere radii (shown in legend). **d**, Angles between the outermost microtubule bundles at the pole in two directions, *α*_*y*_ and *α*_*z*_, (see schemes, *k* = 100 pN/µm) versus compression coefficient. **e**, Spindles before, during and after compression in live-imaged U2OS cells expressing CENP-A (color-coded for depth) and α-tubulin (MTs, gray) with labelled DNA (cyan) in control (top) and mitosis with an unreplicated genome (MUG, bottom) cells. **f**, Setup for spindle compression. **g**, Spindle height (gray) and spindle width (cyan) before, during and after compression for control (top) and MUG (bottom) cells. **h**, Angle between pole and outermost kinetochore fiber for three time points related to compression of U2OS control cells. Thick lines, mean; shaded areas, standard error of the mean (SEM); thin lines, individual cells. Number of cells: 12 (**g**, top and **h**); 11 (**g**, bottom). Abbreviations: Unrepl., unreplicated; MTs, microtubules. The number of cells equals the number of independent experiments. **g**, **h** Student’s t-test. In (**a**), (**b**), (**c**) and (**d**) parameters are as in Fig. 3, unless stated otherwise. Scale bar, 5 µm.

To test the model, we acutely squeezed human RPE1 and U2OS cells in metaphase by compressing them with a gel^66^ (Fig. 4e, f). If the chromosomes indeed push each other as in the model, the metaphase plate is expected to widen, even though different outcomes are possible as there are materials that do not widen under pressure, such as cork, or even become narrower^67^. We found that after 1 minute of compression, spindle height in U2OS cells decreased by ∼44%, the metaphase plate widened by ∼30% and thickened by ∼50%, while spindle length remained unchanged (Fig. 4g, Extended Data Fig. 7c-f, and Movie 2). Similar trends were observed in RPE1 cells (Extended Data Fig. 7c-h). Acute compression also resulted in a wider angle between the outermost microtubule bundles at the spindle pole (Fig. 4h, Extended Data Fig. 7i). When compression was relieved, the spindle and metaphase plate partially regained their original widths in both cell types (Extended Data Fig. 7c), similar to spindles in previous studies^66,68^, irrespective of the extent of height reduction, which ranged from 2 to 12 µm (Extended Data Fig. 7d). Spindles in MUG cells also widened upon compression, but only by ∼23% (Fig. 4e, g), whereas spindle length remained unchanged (Extended data Fig. 7f, and Extended Data Movie 2). Both experiments agree with the model’s predictions that the spindle widens under compression (compare Fig. 4c, d with Fig. 4g, h). For a quantitative comparison, we identified a compression coefficient of 100 pN/μm, where the decrease in spindle height predicted by the model is similar to that observed in experiments. The model suggests that the inter-chromosome pushing forces increased 3-fold in the compression experiments (Fig. 4b). As in experiments, the model predicts a larger spindle widening for the parameters corresponding to control than to MUG cells, 39% and 34%, respectively (Fig. 4a, c). Thus, the biomechanical tests inspired and interpreted by the model confirmed the chromosome crowding hypothesis.

Altered metaphase plate shape caused by acute compression might affect mitotic efficiency. Indeed, metaphase was substantially longer in acutely compressed than in uncompressed RPE1 cells, lasting more than 20 minutes (Extended Data Fig. 7j), compared to less than 10 minutes^69^, respectively. Although extended mitotic duration could result from physical damage to chromosome attachments during compression, this effect is likely minimal, as the majority of compressed cells eventually progressed to anaphase (Extended Data Fig. 7j). Based on these results, we conclude that substantially increased inter-chromosome pushing forces impede mitosis.

## Spindle length and width are independently regulated by different mechanisms

To complement our results on external compression forces, we asked whether endogenous forces, such as cytoplasmic crowding^70^, contribute to maintaining spindle and metaphase plate dimensions. We applied hyperosmotic and hypoosmotic solutions to RPE1 cells in metaphase, which have been shown to increase or decrease cytoplasmic density by approximately 45%, respectively^71^ (Fig. 5a, Extended Data Movie 4). Hyperosmotic shock caused an immediate narrowing of the spindle (∼14%) and the metaphase plate (∼18%), followed by gradual but extensive spindle elongation (∼60%, Fig. 5a, b, Extended Data Fig. 6c, g). These results indicate that increased cytoplasmic forces acutely compress the metaphase plate. In contrast, hypoosmotic shock had no immediate effect but led to gradual shortening of the spindle and widening of both the spindle and metaphase plate (Fig. 5a, b, Extended Data Fig. 6c, g). This finding shows that decreased cytoplasmic density triggers slower remodeling of the spindle and metaphase plate, implying that cytoplasmic pushing forces on chromosomes in unperturbed cells are relatively modest.

**Fig. 5.**
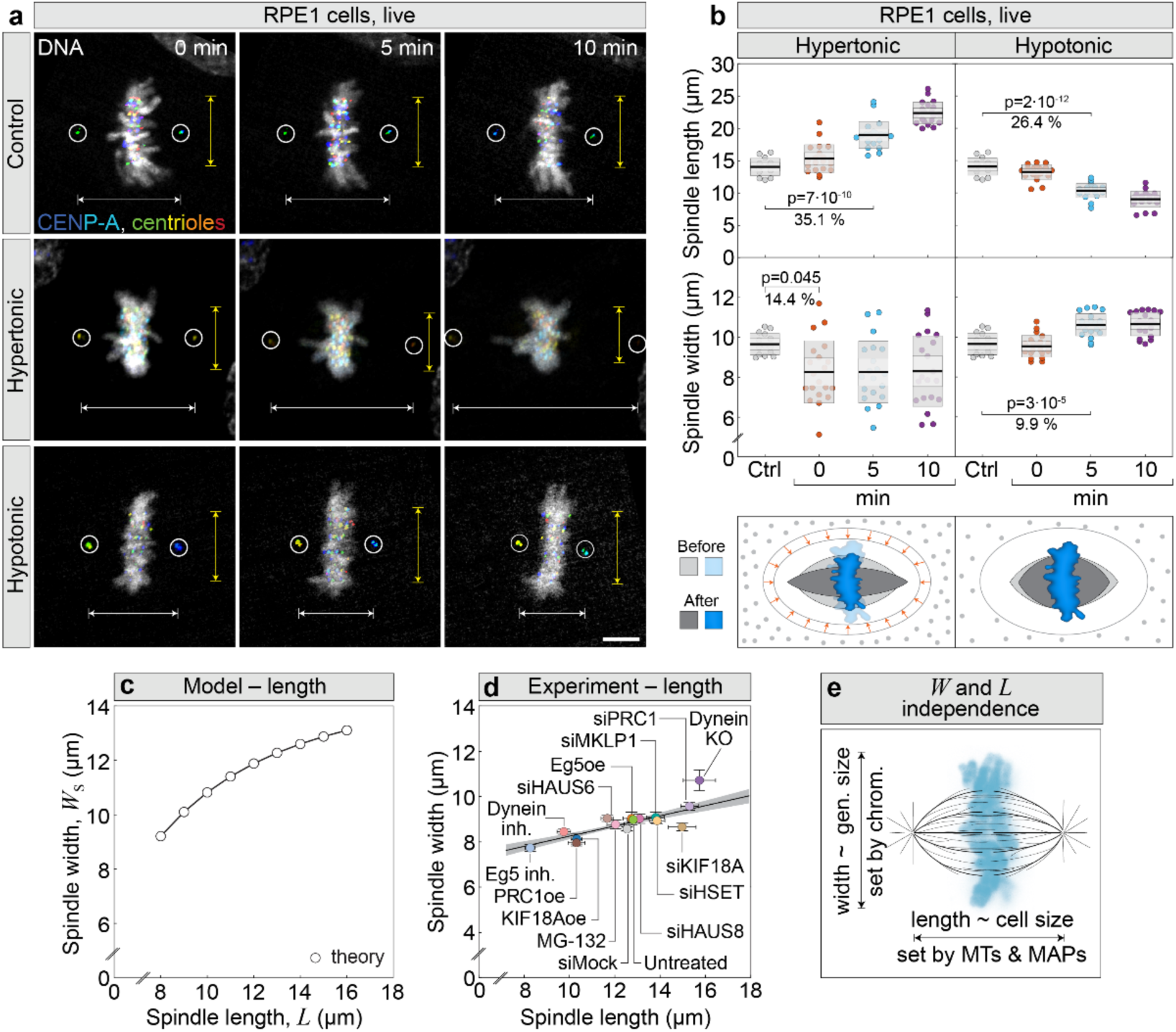
Spindle length and width are independently regulated. **a**, Live-imaged RPE1 cells expressing CENP-A-GFP and centrin-3-GFP (color coded for depth with circled centrioles), and stained with SPY650-DNA (cyan) in control untreated (top), treated with hypertonic solution (middle), and treated with hypotonic solution (bottom). Times denote the start of the imaging. Yellow arrows represent spindle width and white arrows spindle length. **b**, Spindle length (top) and spindle width (bottom) in control (Ctrl) and at indicated times after the addition of hypertonic (left) and hypotonic (right) solution. Colored points represent individual cells; black lines show the mean, with light and dark gray areas marking 95% confidence intervals for the mean and standard deviation, respectively. Number of cells: 16, 27, and 20 per treatment. Percentages indicate differences between mean values of groups. Scheme of spindle and acute cellular responses to hypertonic (bottom left) and hypotonic (bottom right) solutions. Control spindles are shown in lighter colors, and treated in darker colors (legend, left). Orange arrows indicate cell shrinking in hypertonic solution relative to control. Gray dots represent extracellular solute. **c**, Model prediction of spindle width, *W*_*s*_ = *W* − 2*R*, for different spindle lengths. Other parameters are as in Fig. 3. **d**, Mean ± SEM for spindle width versus spindle length for protein perturbation experiments. Data from^72^. Black line, linear fit (R^2^ = 0.58, p = 1.4× *10*^−*25*^); gray area, 95% confidence interval. **e**, Spindle width is determined by the chromosomes and scales with their size. Spindle length is set by microtubules and microtubule-associated proteins (MAPs), and scales with the cell size. In (**c**) parameters are as in Fig. 3, unless stated otherwise. Abbreviations: MTs, microtubules; MAPs, microtubule-associated proteins; gen., genome; inh., inhibited; KO, knock-out; oe, overexpression; si, small interfering RNA; Ctrl., Control. **b**, data are mean with 95% confidence intervals and standard deviation. **b**, Tukey’s Honest Significant Difference test **d**, Wald test. Scale bar, 5 µm.

Acute microtubule depolymerization (Extended Data Fig. 6a, b), external compression (Extended Data Fig. 7k), and osmotic perturbations (Fig. 5a, b) indicate that spindle length and width are regulated independently, because large changes in one dimension were decoupled from the changes in the other. Consistent with the model prediction that a ∼100% increase in spindle length yields only a ∼30% increase in width (Fig. 5c, Extended Data Fig. 7l), perturbation of dozens of mitotic MAPs that regulate microtubule dynamics, sliding, and cross-linking in RPE1 cells^72^, produced a 100% change in length but only a 21% change in spindle width (Fig. 5d). Similarly, spindle length varied with centrosome number at a given ploidy in RPE1 cells, whereas spindle width showed minimal change (Extended Data Fig. 3b, c). Interestingly, mitotic MAPs were dysregulated in adapted tetraploid RPT cells (Extended Data Fig. 4d), suggesting that these alterations contributed to changes in spindle length in these cells. Altogether, spindle length and width are largely independent, indicating that they are indeed controlled by different mechanisms based on regulation of microtubule dynamics^17,73^ and inter-chromosome pushing forces, respectively (Fig. 5e).

## Universality of the chromosome crowding mechanism

As the ultimate aim of our study, we asked whether the proposed mechanism based on inter-chromosome pushing forces is universal and thus can explain the spindle scaling law found across eukaryotes (Fig. 1). To describe different species, we use the model for human spindles and vary only the genome size by modifying chromosome volume, to cover almost 4 orders of magnitude (Box 1). We assume that the volume of a chromosome is proportional to its number of base pairs, as observed for chromosome volume across vertebrate and flowering plant species^28^, and in agreement with a comparison of human and yeast chromosomes^74^ (Methods). Thus, chromosomes of all species are represented in the model by spheres with a radius proportional to the 3^rd^ root of genome size (Box 1). Starting from the experimental data point for human spindles, we calculate the metaphase plate width for both smaller and larger genome sizes, thereby extrapolating the model to regions far beyond its original scope. Remarkably, the model predicts a power-law scaling of the metaphase plate width with the genome size, with an exponent of 0.37, close to 0.33 from experimental data (Fig. 1d, compare dashed and solid lines). Thus, our model together with experiments demonstrates the universality of the chromosome crowding hypothesis across very different eukaryotic species.

The scaling predicted by the model is robust to changes in the number and physical properties of microtubules and chromosomes (Extended Data Fig. 8e; Methods). Moreover, the scaling does not depend on chromosome shape: incorporating a more realistic prolate spheroidal chromosome geometry produces similar scaling behavior (Extended Data Fig. 8e; Methods). In contrast, the scaling directly depends on the relationship between genome size and chromosome volume. When this relationship is nonlinear, the scaling exponent changes markedly (Extended Data Fig. 8f). The model therefore suggests that the experimentally observed scaling exponent of 0.33 arises from a linear relationship between chromosome volume and genome size.

## Discussion

In this work, we identified a conserved mechanism of mitotic spindle adaptation to greatly varying genome sizes from yeasts to plants and animals (Fig. 6). The mechanism is based on metaphase plate and spindle widening driven by inter-chromosome pushing forces. The identified scaling takes into account that chromosome volume is proportional to its amount of chromatin (Fig. 1), which was observed across vertebrates and plants^28^ as well as for individual human chromosomes^29^. We propose a broader principle where the degree of chromatin compaction within the metaphase plate is conserved across diverse evolutionary lineages, even though genome sizes differ by up to four orders of magnitude. The uniform DNA density of metaphase plates across eukaryotes is likely linked to conserved packing machinery, including histones, cohesins, condensins, and topoisomerases^75–77^. However, how this machinery produces mitotic chromosomes of similar density and uniform arm length-to-width ratios across eukaryotes, despite vast genome size variation, remains an exciting topic for future studies.

**Fig. 6.**
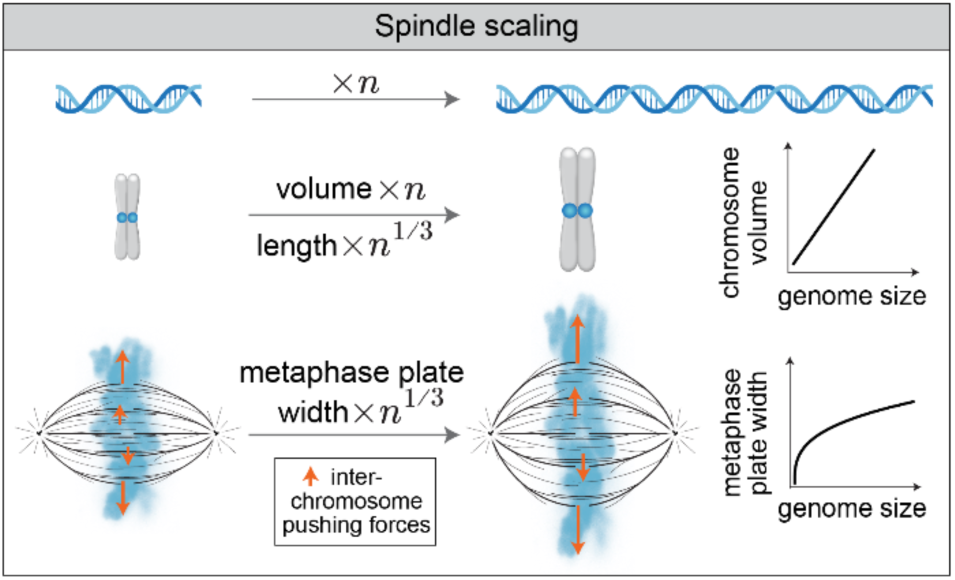
Chromosome crowding in the metaphase plate sets the scaling of the mitotic spindle with the genome size. When DNA length increases *n* times, the volume of a typical chromosome also increases *n* times, while its length and the metaphase plate width increase only *n*^1/3^ times. Thus, a significant change in genome size requires only a small variation in spindle size. The mechanism of spindle adaptation to varying genome sizes is based on spindle widening due to inter-chromosome pushing forces (arrows).

We propose that the magnitude of inter-chromosome forces is influenced not only by chromatin density but also by external forces acting on a cell. This conclusion is supported by our experiments where acute external compression deformed both the cell and spindle, for which the model predicts an increase in inter-chromosome pushing forces from 100 pN to 300 pN (Fig. 4b). This force under compression becomes comparable to the tension force at the kinetochore^62–64^, which may explain prolonged metaphase in our experiments. Previous studies have shown that cells entering mitosis under confinement form disorganized spindles with high rates of mitotic errors^78–80^. Our results suggest that errors under confinement arise from chromosome crowding and excessive inter-chromosome pushing forces, which the spindle microtubules are unable to counteract. The chromosome crowding hypothesis thus explains why cell rounding is a hallmark of eukaryotic cells without a rigid cell wall^81^ and why eukaryotes with larger genomes undergo open mitosis without a nuclear envelope^82^, as both adaptations create more space for the spindle and chromosomes. Moreover, our data indicate that excessive cytoplasmic crowding can exert compressive forces on the metaphase plate and spindle, producing atypical spindle morphologies. This may explain why physiological osmotic balance is maintained within narrow limits during mitosis^83^. The slower osmotic effects on spindle length likely reflect altered microtubule dynamics and tubulin availability^84,85^, while the gradual changes in spindle width and metaphase plate dimensions may reflect alterations in chromosome condensation^71^.

What enables the spindle to reshape itself when faced with an acute increase in inter-chromosome pushing forces? Compression-induced spindle widening occurs even in taxol-treated spindles, suggesting it is independent of microtubule dynamics^66^. Upon external compression, the angle between microtubules at the spindle pole increases (Fig. 4d, h), suggesting that the microtubules pivot around the pole^86–88^. We propose that the process of microtubule pivoting facilitates spindle adaptation in cells with centrosomes. Conversely, in spindles with unfocused poles, such as those in plant cells, microtubules appear straight, indicating smaller compression forces. We speculate that, due to these weaker forces, spindles with unfocused poles can more readily accommodate higher ploidy, facilitating polyploidization as an evolutionary advantage for plants in generating new species.

Chromosome crowding leads to spindle scaling with the amount of chromatin (Figs. 2 and 3b), which explains wider spindles in cells with higher ploidy not only in somatic cells, but also in mammalian oocytes and early embryos. However, during embryogenesis the spindles become narrower even though the number of chromosomes is constant^18,21,89^, likely due to reduction of chromosome size caused by increased chromosome condensation^18,90^. Taking into account that excessive inter-chromosome pushing forces arise under confinement, we propose that increased chromosome condensation is important to control these forces and thereby spindle shape, promoting success of early embryogenesis^91,92^.

Our work reveals that different dimensions of spindle geometry are regulated by distinct mechanisms. Whereas spindle width is determined mainly by chromosome volume, spindle length is controlled by MAPs and influenced by centrosomes in human cells (Figs. 2 and 5e). We propose that these two spindle dimensions are largely independent. This reasoning is in agreement with previous findings that polyploidization in yeast increases spindle width without altering length^93^, and acute microtubule depolymerization in *Xenopus* egg extracts shortens spindles without changing their width^94^. The independent regulation of spindle length and width may promote spindle adaptation to diverse biological contexts by modifying its shape while preserving functionality. Such shape adaptability is likely essential for the proliferation of tetraploid cells, including those naturally occurring in the human liver^95^, and may similarly facilitate the growth of cancer cells that have undergone tetraploidization or become highly aneuploid due to prior chromosomal missegregation^96^. As we found that adapted cells and their spindles are often larger than freshly generated cells of the same increased ploidy (Fig. 2), and this larger size correlates with a distinct transcriptomic signature (Extended Data Fig. 4), alterations in cell size may represent a general mechanism of cellular adaptation to polyploidization^97^.

Metaphase plate thickness, reflecting chromosome alignment, also changed upon polyploidization. We speculate that this arises from altered resources available per chromosome for attachment to spindle microtubules, including kinetochore proteins and MAPs regulating microtubule nucleation and dynamics. Our transcriptome analysis indicates that the mitotic transcriptome increases sublinearly with ploidy^46^, implying fewer mitotic factors per chromosome. These changes may weaken microtubule attachments, enhance kinetochore oscillations, and overall impair chromosome alignment following polyploidization.

Taking a broader perspective across all eukaryotes, we propose that the conserved packing of mitotic chromosomes requires a rather small variation in spindle size for significant changes in genome size. These features together allow the spindle to be built upon the same biological principles across eukaryotes, helping cell diversification without a need for new cell division mechanisms. We suggest that the spindle’s biomechanical adaptability promotes evolution by allowing the division of substantially altered genomes after speciation-driving events such as polyploidization^98^.

## Methods

### The model for mitotic spindle based on the inter-chromosome pushing forces

#### Geometry

In our three-dimensional model, the mitotic spindle has length *L* and it is composed of *N* microtubule bundles that extend between two spindle poles, where the geometry is parametrized by Cartesian coordinates with *x*-, *y*-, and *z*-axes, and the associated unit vectors, **x̑**, **ŷ**, and **z̑** (Fig. 3, Extended Data Fig. 8a-d). In our description, the spindle is aligned to the *x*-axis, with the left and right poles located at points (0, 0, 0) and (*L*, 0, 0), respectively (Extended Data Fig. 8a). Poles are connected by the radius vector **L** = *L***x̑**. A point along the contour of bundle *i* is denoted by radius vector **r**_*i*_(*S*), which is a function of contour length *S* (Extended Data Fig. 8d). In order to simplify calculations, we use a small angle approximation, *S* ≈ *x*, which is a plausible approach for the majority of spindle microtubules^99^. A chromosome is bound to microtubule bundle at position **r**_*i*,chr_, with projection onto *x*-axis that is equal for all chromosomes, *x*_chr_ = **r**_*i*,chr_ ⋅ **x̑**. In the case of chromosomes aligned within the metaphase plane they obey *x*_chr_ = *L*/2 (Extended Data Fig. 8d). Positions of the microtubule bundle ends, **d**_*i*,left_ ≡ **r**_*i*_(0) and **d**_*i*,right_ ≡ **r**_*i*_(*L*) − **L**, are defined relative to the left and right poles, respectively.

#### Mechanical balance

In our theory we consider the mitotic spindle as a static object. Thus, we impose a balance of forces and torques for each individual bundle and pole. The balance of forces for the bundle *i* reads

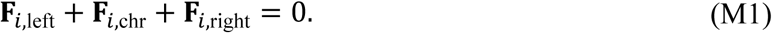

Here, vectors **F**_*i*,left_ and **F**_*i*,right_ denote forces exerted on the bundle by the left and the right pole, respectively. We also consider force **F**_*i*,chr_ which is exerted by the chromosome attached to bundle *i*. This force arises from inter-chromosome pushing forces,

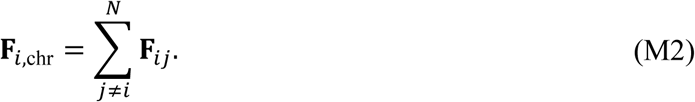

Here, vector **F**_*i*j_ denotes the force by which chromosomes *j* pushes chromosomes *i*, with the opposing force obeying **F**_j*i*_ = −**F**_*i*j_(Extended Data Fig. 8c). In Box 1, Eq. (3) offers a simplified version of Eq. (M2), explicitly stating in words that the sum is restricted to contributions from neighboring chromosomes. The torques for the bundle *i* are also in balance, which for the left bundle end reads

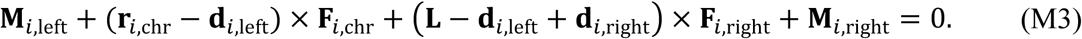

Here **M**_*i*,left_ and **M**_*i*,right_ denote torques exerted by two respective spindle poles on the bundle *i*. Eqs. (M1)-(M3) together describe the mechanical balance of the individual microtubule bundles.

The model also includes balance for the spindle poles. We first introduce a theoretical description for the left pole. Balance of forces and torques for the left pole read:

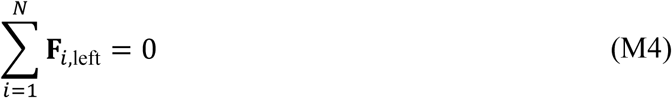

and

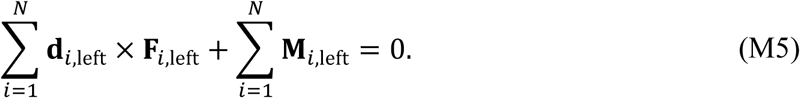

Experiments show that microtubule bundle pivots around the spindle pole under external force^87^, which we include in the model by assuming a freely joint condition **M**_*i*,left_ = **M**_*i*,right_ = 0. Note that balance at the right pole is described by an analogous set of equations which can be obtained by replacing index ‘left’ with index ‘right’ in Eqs. (M4) and (M5). These equations, together with Eqs. (M1)-(M3), describe the mechanical balance of the entire spindle.

#### Inter-chromosome pushing forces

A chromosome is modeled as an elastic sphere, whereas in the next step we generalize chromosome geometry by modeling it as a prolate spheroid. Spheres are parameterized by the radius *R* and the corresponding volume, 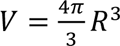. The genome size is proportional to total volume of all chromosomes, *G* = *ρNV*, where *ρ* denotes the DNA density^28^. This relationship is given in Box 1 as Eq. (2). Thus, a direct link between sphere radius and genome size is given by

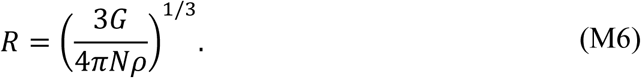

A chromosome deforms under pushing forces exerted by its neighbors. The reduction in the distance between chromosomes, ℎ_*i*j_ = (2*R* − |**r**_*i*,chr_ − **r**_j,chr_|)*θ*(2*R* − |**r**_*i*,chr_ − **r**_j,chr_|) (Extended Data Fig. 8c), reflects the extent of their deformation (Fig. 3a). Here, the Heaviside step function *θ* ensures that there is no force acting on spheres if they are not in contact. This deformation results with repulsive force, which for small deformations reads^100^

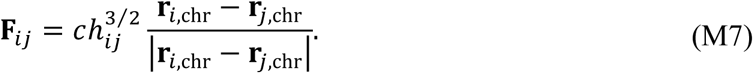

A simplified version of Eq. (M7) is given as Eq. (1) in Box 1, where we calculate a magnitude of inter-chromosome pushing force only. The elasticity constant *c* is calculated assuming that spheres are elastic objects described by Young’s modulus *E* and Poisson ratio *σ*,

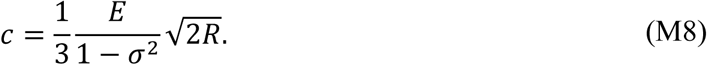

This description of inter-chromosome pushing forces considers elastic properties of homogenous spheres and is valid for small deformations, which is the case in our system (Fig. 3a, d). We do not consider short-range electrostatic surface interactions because previous experiments have shown that they have a small effect on the metaphase plate width^36^. Eqs. (M6)-(M8) describe pushing forces between two chromosomes modeled as spheres with a given genome size and DNA density.

In order to be closer to chromosome shape, we generalize its geometry by modeling them as prolate spheroids with the polar radius *R*_1_ and the equatorial radius *R*_2_. In this case, the inter-chromosome pushing force is also given by Eq. (M7), whereas the elasticity constant changes to^101^

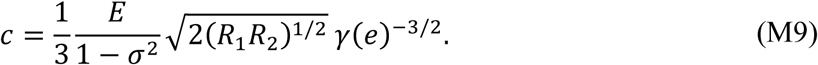

For a special case when polar and equatorial radii are equal, *R* = *R*_1_ = *R*_2_, the prolate spheroid becomes a sphere and Eq. (M9) reduces to Eq. (M8). The auxiliary function in Eq. (M9) depends on the eccentricity of the contact ellipse, *e*, and is given by

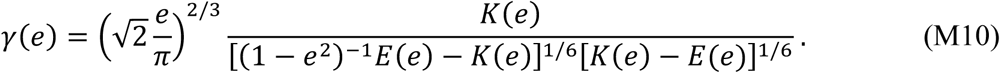

Here, the elliptic integrals of the first and the second kind are given as 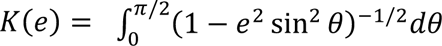 and 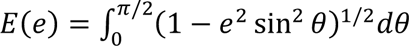, respectively. The eccentricity of the contact region ellipse is given implicitly by

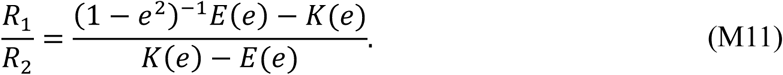

In our calculations, we evaluate *e* by numerically solving this equation.

#### Shape of flexible microtubule bundle

In order to predict the shape of microtubule bundles, which sets the geometry of the spindle, we calculate how microtubules deform under force and torque. We parametrize microtubule bundle shape by the unit tangent vector, **t**_*i*_(*S*) = *d* **r**_*i*_⁄*dS* (Extended Data Fig. 8d), and the angle that describes the orientation of the cross-section relative to one of the straight bundle, *Φ*_*i*_(*S*). The shape of the slender rod is given by the static Kirchhoff equation,

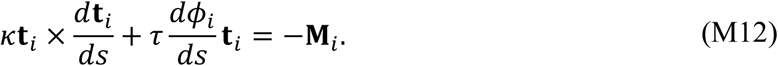

Here, the parameters *κ* and *τ* denote flexural and torsional rigidity, respectively. To calculate the torque in Eq. (M13), we solve the balance of torque on segment of elastic rod,

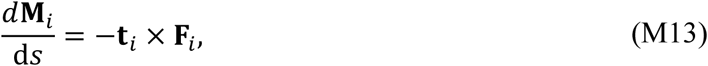

where **F**_*i*_ denotes a contact force at position *S*. To integrate Eq. (M13) we take into account that contact force has a jump at *x* = *L*⁄2. In the small angle approximation we obtain **M**_*i*_(*x*) = **M**_*i*_(0) − (**r**_*i*_ − **d**_*i*,left_) × **F**_*i*,left_ for *x* < *L*⁄2 and **M**_*i*_(*x*) = **M**_*i*_(0) − (**r**_*i*_ − **d**_*i*,left_) × **F**_*i*,left_ − (**r**_*i*_ − **r**_*i*,chr_) × **F**_*i*,chr_ for *x* > *L*⁄2. After implementing boundary conditions **M**_*i*_(0) = **M**_*i*_(*L*) = 0 the contact torque along the contour, exerted by the portion of rod from the left microtubule end, is calculated as

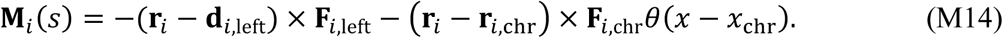

The Heaviside step function ensures that force **F**_*i*,chr_ is exerted at the position of chromosome *i*. We also assume that microtubule bundles have the same force parallel with the main spindle axis and from Eq. (M4) follows that it has a vanishing value, **x̑** ⋅ **F**_*i*,left_ = 0.

In order to solve the model, we first calculate a microtubule shape by solving Eq. (M12). We solve it in a small angle approximation, together with the other previously stated assumptions, namely (i) chromosomes are positioned within the metaphase plane, (ii) the microtubule-pole connection is freely jointed, and (iii) there is no force parallel with the main spindle axis. In the case of a spindle with mirror symmetry, where the metaphase plate is a plane of symmetry, Eqs. (M1) and (M3) yield 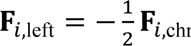. In this case the corresponding microtubule bundle lies within a plane. In particular, for force exerted by the chromosome pointing in the direction of *y* axis and magnitude *F*_*i*,chr_ ≡ |**F**_*i*,chr_|, Eq. (M12) simplifies to *d*^2^*y*/*dx*^2^ = −*F*_*i*,chr_*x*/(2*κ*), for *x* < *L*/2, which is given as Eq. (4) in Box 1.

#### Position of microtubule bundle ends

Microtubule bundle ends are located at nodes of the finite two-dimensional hexagonal lattice (Extended Data Fig. 8b). The lattice parameter, *α*_0_, represents the nearest-neighbor distance between lattice nodes situated within a single plane. We consider the cases where *N* = 1 + 3(*m* + 1)*m* nodes are distributed within the lattice such that comprise a regular hexagon with edge of length *α*_0_*m*. For this reason, we consider the cases with *N* = 7, 19, 37, 61, 91 microtubule bundles which correspond to *m* = 1, 2, 3, 4, 5, respectively. The center of the lattice corresponds to spindle pole, whereas the lattice plane is perpendicular to **x**-axis.

#### External compressive force

To describe experiments in which spindles are compressed by external forces, we change the Eq. (M2) by adding the force, **F**_*i*,ext_, that arises from interactions of chromosomes with the external probe,

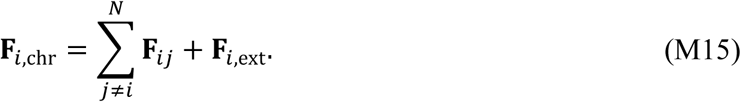

Interactions with the external probe are approximated by the linear elastic restoring force, **F**_*i*,ext_ = −*k*(ŷ ⋅ **r**_*i*,chr_)ŷ, where *k* denotes a phenomenological compression coefficient. In experiments, external forces influence the spindle poles, ensuring the spindle remains aligned with the cover slip. In our theoretical approach, this is modeled by constraining the spindle pole positions to remain located at points (0, 0, 0) and (*L*, 0, 0) under compression.

#### Numerical procedure

For numerical computations of the complete spindle, we reformulate the model as a problem of minimizing the total energy. Under the small-angle approximation, the potential bending energy of the *i*th bundle, as described by Kirchhoff’s equation (M12), is expressed as

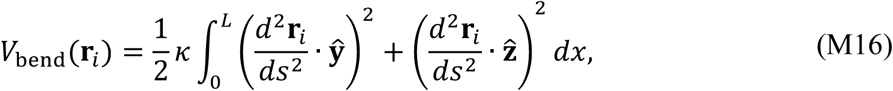

where **r**_*i*_ · ŷ and **r**_*i*_ · **z̑** describe transversal displacements. For a detailed derivation of these equations from three-dimensional linearized elasticity, refer to^102^. The potential associated with the repulsive force between *i*th and *j*th chromosomes, as given in Eq. (M7), is given by

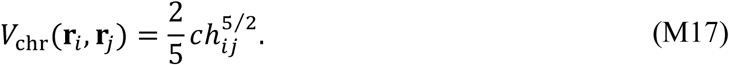

The third potential we consider arises from the linear elastic restoring force at the *i*th chromosome. This potential is given by

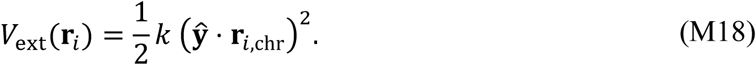

The total potential of the entire spindle, incorporating the repulsive forces between chromosomes and the restoring forces, can now be expressed as the sum of all potentials from Eqs. (M16), (M17), and (M18) across all bundles with parametrizations collected in **r** = (**r**_1_,…, **r**_*N*_)

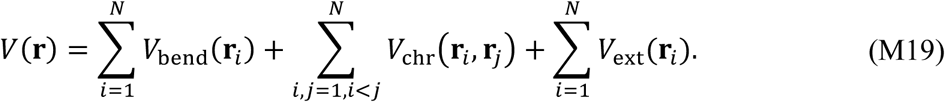

Then the equilibrium configuration **r** of the spindle minimizes the functional *V* over the set of bundle parametrizations that satisfy the specified boundary conditions. To solve this minimization problem, we discretize it using the finite element method as described in^103^ and employ the gradient descent type method as in^104^ to approximate the minimizer. Our model for the mitotic spindle is given by Eqs. (M1)-(M19). However, for understanding basic ideas behind the inter-chromosome pushing hypothesis and its relevance for spindle mechanics, we provide its simplified version in Box 1.

#### Stability analysis of the obtained spindle shape

In our model, chromosomes are constrained to align at the metaphase plane, whereas in living cells they can be displaced from it. We therefore extend the model by introducing a force directed toward the metaphase plane that still allows such displacements. We perform a stability analysis for the equilibrium solution by calculating the second variation of the energy functional and evaluating under which condition it has a positive value.

To describe a centering force that positions chromosomes, we extend the energy functional in Eq. (M19) by adding an extra term,

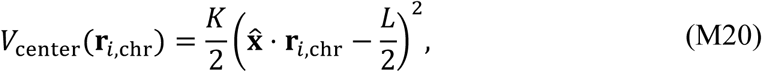

which describes a harmonic centering potential. Here, parameter *K* denotes strength of centering potential. To calculate the second variation of the energy functional, we consider variation of shapes of microtubule bundles, as well as the variation of spheres’ positions. We add a variation to the unperturbed microtubule bundle shape, **r**_*i*_(*x*) + *ϵη*_*y*,*i*_(*x*)ŷ + *ϵη*_*z*,*i*_(*x*)**z̑**. Here, *η*_*y*,*i*_ and *η*_*z*,*i*_ are arbitrary functions whereas *ϵ* is any number close to zero. We also add a variation to the unperturbed chromosome position, 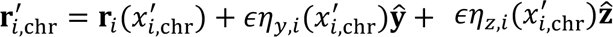, where 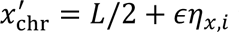 is its perturbed *x*-coordinate.

We expand the position of the spheres in Taylor series up to the second order, since this is the highest order contributing to the second variation of the energy functional. We obtain

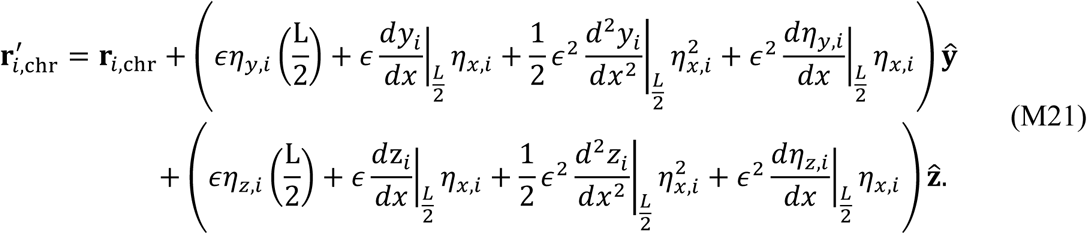

For the variation of the microtubule bundle shape we assume that it is described by functions which obey conditions *η*_*y*,*i*_(*L*/2) = 0 and *η*_*z*,*i*_(*L*/2) = 0. This allows us to obtain an asymmetric microtubule bundle shape for chromosomes displaced from the metaphase plane.

The second order variations of Eq. (M19), together with centering potential Eq. (M20), will result with changes in three terms, representing microtubule bending, inter-chromosome pushing forces, and centering force, 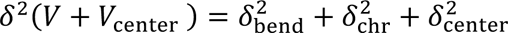. The respective terms are given by:

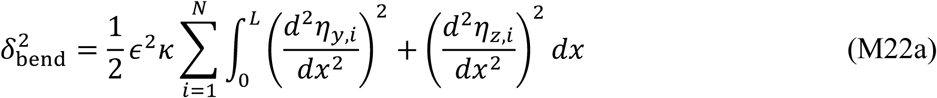

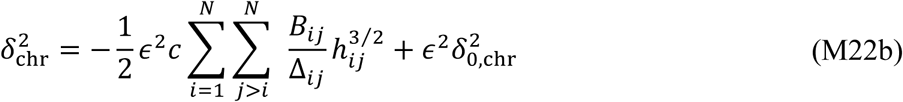

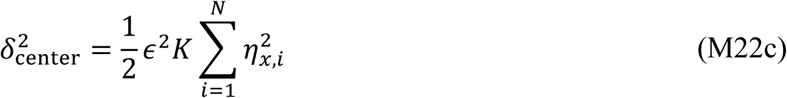

In the second variation of energy, displacement from the metaphase plane can appear in the case of possibly negative first term in Eq. (M22b). Thus, we decided to write this equation by separating the term with negative value from the term 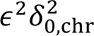 with positive value. We also use the shorthands *Δ*_*i*j_ ≡ |**r**_*i*_ − **r**_*j*_|, and

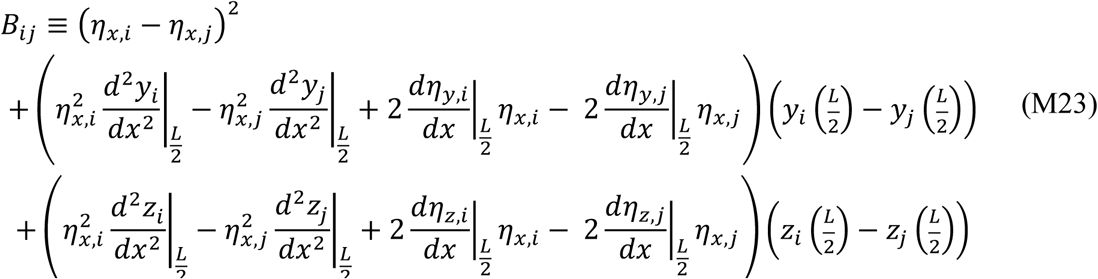

Because the second variation of energy functional of our system results with a very complex solution, we decided to find a lower limit on centering spring and flexural rigidity and compare them with biologically relevant values. We estimate the lower bound in Eq. (M23) by using Young’s inequalities,

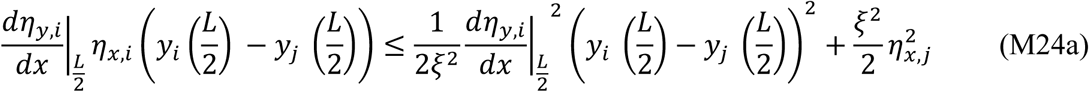

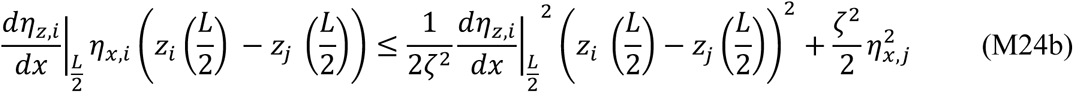

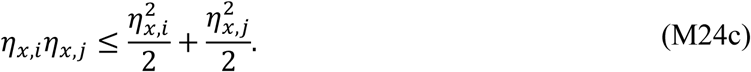

These inequalities hold for any choice of parameters *ξ*, *ζ* ≠ 0. Further, we apply the Poincaré inequality for the function with fixed ends^105^,

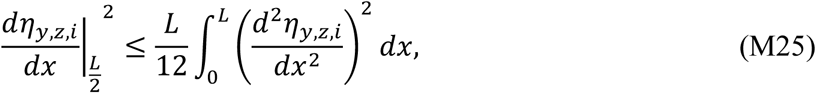

which is used to calculate Eqs. (M24a) and (M24b). Further, we combine Eqs. (M22), (M23), and (M24) which results in a lower bound of the second variation of energy functional. We calculate in which case the sum of the factors that multiply 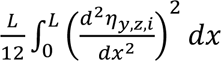 and 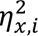, and appear in both 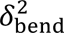 and negative term in 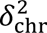, vanishes. In the first case, we obtain an estimate for lower bound on flexural rigidity,

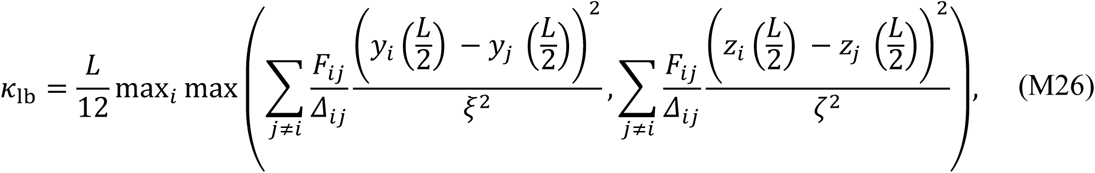

whereas in the second case we obtain an estimate for lower bound on strength of centering potential,

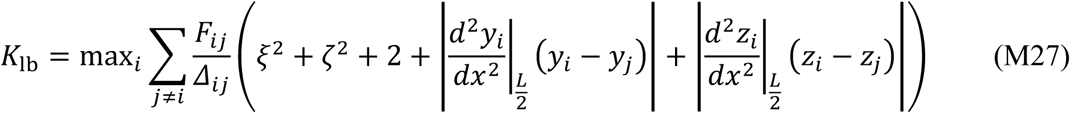

In Eqs. (M26) and (M27) we define *F*_*i*j_ = N**F**_*i*j_N. By evaluating Eqs. (M26) and (M27) we obtain a lower limit for microtubule flexural rigidity and centering potential which we use to calculate stability of the metaphase plate.

To determine the stability conditions for chromosomes aligned in the metaphase plane, we estimate lower bounds on the flexural rigidity and the centering potential strength by evaluating Eqs. (M26) and (M27) using parameters taken from Fig. 3. The central sphere experiences the largest inter-chromosome pushing forces from its neighbors, with *F*_*i*j_ ≈ 50 pN (see Fig. 3d). Because the spheres are only slightly deformed, the distance between their centers can be approximated as twice the radius, *Δ*_*i*j_ ≈ 2.5 µm. Furthermore, the curvature of the central bundle vanishes 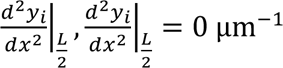. For the arbitrary constants in Eqs. (M26) and (M27), we choose *ξ* = 1 and *ζ* = 1. For this choice of parameters, lower bounds on flexural rigidity and centering potential strength are *k*_lb_ = 450 pNµm^2^ and *K*_lb_ = 480 pNµm^−1^, respectively. The value of flexural rigidity parameter for microtubule bundles is calculated from value for a single microtubule, *κ* = 900 pN · µm^2 106,107^. This value is indeed above the lower bound *k* > *k*_lb_. Centering potential strength, however, is not measured directly and thus we estimate its value from measured kinetochore forces and displacement from the metaphase plate. Owing that forces acting on chromosomes are several hundreds of pN^62–64^ and typical displacement of chromosome is about 1 µm^108^, estimated centering potential strength is also several hundreds of pN/µm and thus comparable with lower bound *K*_lb_.

#### Choice of parameters

In our model we use 8 parameters, which are either estimated from literature or determined in this study. The Young modulus, *E* = 400 Pa, is taken from^109^, where it corresponds to the extension measurements of mouse embryonic fibroblasts in mitosis. Poisson ratio, *σ* = 0.3, is taken as a typical value in^61^ and corresponds to the extension measurements of newt lung cells in mitosis. DNA density, *ρ* = 36 Mbp/μm^3^ = 0.037 pg/μm^3^ ⋅ 978 Mbp/pg, has been used to reproduce metaphase plate width for human cells in Fig. 1d. Spindle length for human cells, *L* = 14 µm has been measured in this study. Number of the chromosomes is varied among the values *N* = 7, 19, 37, 61, 91, whereas the value *N* = 37 is used in most of the calculations. Sphere radius is varied from 0.5 µm (0.05 µm for Fig. 3b) to 2 µm, whereas the value *R* = 1.25 µm is used in most of the calculations related to human spindles, to reproduce the observed metaphase plate width with *N* = 37. The lattice parameter, *α*_0_ = 0.25 µm is chosen to yield the pole diameter of 1.5 µm for *N* = 37, whereas smaller value, *α*_0_ = 0.0025 µm, is used in calculations presented in Figs. 1d and 3b, which is necessary to access the width of spindles with small chromosomes. The phenomenological compression coefficient is varied from 0 to 1000 pN/µm, and value *k* = 100 pN/µm is chosen in some of the calculations (Fig. 4i, Extended Data Fig. 6a, b and k), to reproduce the observed change in spindle width and height following the compression, with *N* = 37.

### Estimation of the relationship between chromosome size and genome size

To apply the model to different species, we need to consider how chromosome shape, size, and number change with the genome size. We chose to obtain these relationships by comparing yeast and human chromosomes because they are well studied and represent small and large genomes, respectively, differing by a factor of 280.

Mitotic chromosomes have a well-defined X-shape with two chromatids each having two arms, where arm length and width is characteristic to species and individual chromosomes. An average mitotic chromatid arm in the yeast *S. cerevisiae* is 0.12 μm wide and around 0.5 μm long, whereas in healthy human RPE1 cells a chromatid arm is 0.65 μm wide and 3 μm long^74^. Thus, both length and width increase by a factor of 5–6, suggesting isometric scaling of typical chromosomes between yeast to humans. This conclusion can be extended to other species based on similar chromosome shapes visible in karyograms of different organisms. Thus, chromosome size of different species can be described by a characteristic length.

To link the characteristic length of a typical chromosome with the species genome size, we assume that the volume of a chromosome, which scales with the characteristic length, is proportional to its number of base pairs. We justify this assumption by the following back-of-the-envelope calculation. The average human chromosome has 3400 Mbp / 23 = 148 Mbp, as the haploid number of chromosomes is 23, and the average yeast chromosome has 12 Mbp / 16 = 0.75 Mbp, as the haploid number of chromosomes is 16, yielding a ratio of 197. Based on our assumption, an average human chromosome has 197 times larger volume than yeast chromosome, implying that characteristic length of human chromosome is 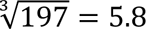 times larger than in yeast. Remarkably, this ratio is within the range of 5–6 obtained from the increase in chromatid arm length and width from yeast to humans described above, supporting our assumption. Thus, the characteristic length of a typical chromosome scales with the 3^rd^ root of genome size.

### Cell lines

The cell lines used in this study were: **1.** human hyper-triploid (modal chromosome number = 74) U2OS (human osteosarcoma, female) cells expressing CENP-A-GFP and mCherry-alpha-tubulin, which were a gift from Helder Maiato (Institute for Molecular Cell Biology, University of Porto, Portugal), **2.** human diploid (modal chromosome number = 46) hTERT-RPE1 (retinal pigmented epithelium, female) cells expressing CENP-A-GFP and centrin1-GFP, which were a gift from Alexey Khodjakov (Wadsworth Center, New York State Department of Health, Albany, NY), **3.** human diploid hTERT-RPE1 cells and their hypotetraploid adapted clones, RPT1, RPT3 and RPT4 (all with modal chromosome number = 80) expressing H2B-GFP, established as described previously^38^, **4.** human diploid hTERT-RPE1 cells with or without p53 knockdown (KD) expressing H2B-Dendra2, which were a gift from Rene Medema (Netherlands Cancer Institute, Amsterdam, Netherlands), **5.** Human near-diploid (modal chromosome number = 45) HCT116 (human colorectal carcinoma cell line, male) cells, and their hypotetraploid adapted clones HPT1 (modal chromosome number = 75) and HPT2 (modal chromosome number = 78), established as described previously^38^.

### Cell lines culturing

Cells were cultured in flasks containing Dulbecco’s modified Eagle’s medium (DMEM; Capricorn Scientific), supplemented with 10% fetal bovine serum (FBS; Sigma-Aldrich) and a 10000 U/mL penicillin/streptomycin solution (Capricorn Scientific). The cultures were maintained at 37°C with 5% CO_2_ in a Galaxy 170S CO_2_ humidified incubator (Eppendorf) and routinely subcultured upon reaching 70–80% confluence. Prior to experimentation, all cell lines were screened for mycoplasma contamination through monthly assessments using the MycoAlert Mycoplasma Detection Kit (Lonza), and additional checks during imaging procedures with DNA labelling dyes.

### Establishment of 3D cultures of mouse primary cells

Mouse small intestine (mSI) organoids were derived from the small intestines of female wild-type C57BL/6 mice, sourced from the Laboratory for Neurodegenerative Disease Research at the Ruđer Bošković Institute, in compliance with the EU Directive 2010/63/EU for animal experimentation and the applicable national regulations of Croatia. The mSI organoids, genetically modified for stable expression of eGFP-centrin-2, were established using small intestines obtained from mice resulting from crossing of transgenic GFP-CETN2 mice (JAX stock #008234, The Jackson Laboratory)^110^ and transgenic Cre-inducible H2B-mCherry mice (JAX stock #0231239, The Jackson Laboratory)^111^.

To isolate small intestine crypts from mice, the small intestine was cleaned in cold PBS and cut in pieces of 2–5 mm. Cleaned pieces were incubated for 1h in cold 20 mM EDTA in PBS while shaking. Supernatant was collected repeatedly, each collection resulting from incubating the intestine pieces in fresh, cold EDTA solution for 5–10 minutes and passing the agitated suspension through a 70 µm cell strainer. The highest quality collection was centrifuged at 300 g for 7 min at 4°C and the pellet was resuspended in cold PBS. Suspension was mixed with culture medium and Matrigel (Corning) was added in a 1:2 ratio. The final mixture was plated in pre-warmed 24-well plates.

Organoid cultures were maintained in IntestiCult™ Organoid Growth Medium (Mouse, STEMCELL Technologies) and passaging was performed using mechanical fragmentation. For experiments, organoids were dissociated by incubation with TrypLE Express (Thermo Fisher Scientific) for 5 min at 37°C. Dissociated single cells and clumps were plated in Matrigel domes on 35-mm uncoated dishes with 0.17 mm glass thickness (MatTek Corporation), covered with 500 µl of the IntestiCult™ medium and cultured for 24–96 h prior to fixation. Samples fixed 24 h post-isolation were cultured with addition of 10 µM Y-27632 (ROCK inhibitor, HY-10583, MedChemExpress) and 1 h before the fixation the inhibitor was washed out.

### Human colorectal cancer organoids

Movies of patient-derived organoids (PDOs), representing various types of colorectal cancer, were partly obtained from a previous study^47^, while additional movies acquired using the same methods were not included in the original publication. The analyzed PDOs included lines established from tumor resections (24Ta, 2Tb, 9T, 7T) and one line derived from healthy tissue adjacent to a tumor (P1N1)^47^. Additionally, the PDO line P27T, established from a tumor resection, was excluded from the original publication, but was cultured and imaged under the same conditions as the other PDO lines^47^.

### RNA interference

RPE1 cells 1 day prior to CENP-E depletion, were seeded onto 35 mm glass coverslip dishes with a glass thickness of 0.17 mm (Ibidi) and allowed to reach 50% confluence through incubation. Small interfering RNA (siRNA) constructs were diluted in Opti-MEM (Gibco), and transfection was carried out using Lipofectamine RNAiMAX Reagent (Invitrogen) as per the manufacturer’s instructions. The ON-TARGETplus Control Pool Non-Targeting pool (D-001810-10-05, Dharmacon), human CENP-E ON-TARGETplus SMART pool siRNA (L-003252-00-0010, Dharmacon), and human TP53 ON-TARGETplus SMART pool siRNA (L-003329-00-0010, Dharmacom) constructs were used at a final concentration of 100 nM. Following a 4-hour incubation period with the transfection mixture, the medium was replaced with fresh cell culture medium. Fixation subsequent to CENP-E depletion was executed 48 hours after transfection. Depletion effectiveness of used CENP-E siRNAs compared with a non-targeting control was previously measured using the same RNA interference protocol, showing near-total depletion of CENP-E^49^.

### Inducing mitosis with an unreplicated genome (MUG)

MUG induction in U2OS cells was achieved through incubation with hydroxyurea and caffeine^51^. Stock solutions of hydroxyurea (HY-B0313, MedChemExpress) and caffeine (C0750, Sigma-Aldrich) were prepared at a concentration of 20 mM each. These stock solutions were diluted in DMEM to obtain final concentrations of 2 mM for hydroxyurea and 5 mM for caffeine. U2OS cells were seeded in 2 mL of DMEM at 37°C with 5% CO_2_ on 35 mm uncoated glass coverslip dishes (MatTek Corporation). After 12 hours, the culture medium was replaced with 1 mL of DMEM containing 2 mM hydroxyurea, and the cells were incubated for an additional 20 hours. Subsequently, the medium was replaced with fresh medium containing 2 mM hydroxyurea and 5 mM caffeine, and the cells were further incubated for 18 hours before imaging for compression experiments, and 13 hours for cell and spindle morphometry experiments due to the larger number of mitotic cells.

### Spindle compression assay

The spindle compression method was refined based on previous work^66^, as described in^72^. A solution comprising 2% ultra-pure agarose (15510, Thermo Fisher Scientific) in PBS was prepared, heated to boiling, and poured into a 35 mm petri dish to solidify, achieving a final thickness of approximately 0.5 cm. During the solidification process, thin strings were inserted into the agarose solution, leaving one end inside the dish while the other remained free outside. Gel areas measuring 1 cm × 1 cm (with strings attached) were excised and stored in PBS at 4°C, then warmed to 37°C immediately before use in experiments. Cells in metaphase were selected from cultures with 80–100% confluence. Following imaging of the metaphase cell before compression, the gel was delicately deposited, centered on the cell. Compression was executed using an oil hydraulic fine manipulator (InjectMan 4, Eppendorf) with a dynamic movement control (100–240 V/50–60 Hz) and a coarse manipulator connected to the confocal Opterra I microscope. A metal rod (part of the micromanipulator) was positioned centrally above the gel, and gradually lowered along the z-axis until gentle contact was made with the gel (rod diameter significantly larger than cell diameter). The rod was then slowly lowered further (over approximately 10 seconds) by several micrometers until the cell area expanded, as evidenced by the appearance of membrane protrusions. Cell and spindle responses were continuously imaged while maintaining the position of the cell. Subsequently, the cells were imaged for a third time immediately after removing the rod and releasing the pressure. This provided three time points: images before compression, during compression (for 1 minute), and shortly after release. In some instances, cells were left to recover for an extended period to assess their ability to divide before being imaged again. Spindles with a height change of less than 20% during compression were excluded from the analysis.

### Induction of mitotic slippage and cytokinesis failure with centrosome elimination

Tetraploid cells were generated from RPE1 cells with p53KD expressing H2B-Dendra2, while hypooctaploid cells were generated from RPT3 cells expressing H2B-GFP. The cells were seeded in 2 mL of DMEM at 37°C with 5% CO_2_ on 35 mm glass coverslip dishes (MatTek Corporation). The following day, the medium was replaced, and fresh DMEM containing the Plk4 inhibitor centrinone^34^ (HY-18682, MedChemExpress) at a final concentration of 300 nM was added for 24 hours. The following day, the medium was replaced again, and fresh DMEM was replaced sequentially five times. To induce mitotic slippage, cells were treated with 100 μM monastrol (Eg5 inhibitor, HY-101071A, MedChemExpress) and 500 nM AZ3146 (Mps1 inhibitor, HY-14710, MedChemExpress) for 24 hours. The next day, DMEM was replaced five times. After 5 hours, samples were either fixed or subjected to live imaging. Freshly thawed centrinone was exchanged twice a day to prevent Plk4 reactivation and centriole over-duplication. The resulting cell population contained both cells with 1:0 and 1:1 spindles (see Extended Data Fig. 2c). To obtain a predominantly population of 1:1 cells, 24 hours after seeding, cells were treated with CDK4/CDK6 inhibitor palbociclib (250 nM, HY-50767, MedChemExpress)^112^. The following day, the medium was replaced sequentially five times with fresh DMEM, and cells were simultaneously treated with centrinone, monastrol and AZ3146, in concentrations described above. After 24 hours, the medium was replaced five times with fresh DMEM, and cells were left for 5 hours before imaging or fixation (see Extended Data Fig. 2c). To improve polyploidization efficiency, we implemented an optimized protocol (see Extended Data Fig. 2c). Thymidine (2 mM, Sigma, T1895) was used for G1 synchronization for 24 h, followed by three washouts in warm DMEM. In RPT3 clones, TP53 was depleted 4 h prior to synchronization using 100 nM siRNA (details described above) to enhance release and promote subsequent division of polyploid cells. Cytokinesis failure was induced by treating cells with cytochalasin D (750 nM) for 20 h. Centrinone (300 nM) was added continuously each day following thymidine washout. After cytochalasin D washout for three times with fresh DMEM, cells were incubated for an additional 40 h to allow progression toward the next division. To enrich cells in metaphase, the proteasome inhibitor MG-132 (5 µM, Merck, M7449) was added 2 h before fixation. DAPI (1 µg/mL) was added 20 min prior to imaging, and concentration was kept constant for all imaged conditions and replicates.

### Induction of cytokinesis failure

Tetraploid RPE1 cells were generated from both p53 wild type and p53 knockdown cells expressing H2B-Dendra. The cells were seeded in 2 mL of DMEM at 37°C with 5% CO_2_ on 35 mm glass coverslip dishes (MatTek Corporation). The following day, the cells were treated with either 0.75 µM Cytochalasin D (inhibitor of actin polymerization, HY-N6682, MedChemExpress) or 50 µM Blebbistatin (inhibitor of non-muscle myosin II., HY-120870, MedChemExpress), and incubated for 18 h overnight. For live cell experiments, the cells were washed four times with 1X PBS and then transferred to the imaging medium. After 6 hours, the cells were taken to the microscope for imaging.

### Nocodazole and osmotic shocks

hTERT RPE1 cells expressing CENP-A-GFP and centrin-GFP were incubated overnight with 20 nM SPY-DNA-650 (Spirochrome) and acutely treated with nocodazole (HY-13520, MedChemExpress, 10 mM stock in DMSO) to a final concentration of 1 µM. Approximately 1 h prior to the start of the imaging cells were treated with 5 μM MG-132. Imaging was done continuously for 20 min post-treatment, with nocodazole added directly on the microscope stage to predefined metaphase cell positions (details in Microscopy section). Parallel samples were fixed at 0, 3, and 7 min after nocodazole addition and immunostained for α-tubulin to confirm microtubule depolymerization (details in Immunoflorescence section).

hTERT RPE1 cells expressing CENP-A-GFP and centrin-GFP were incubated overnight with 20 nM SPY-DNA-650 (Spirochrome). Approximately 1 h hour prior to the start of the imaging cells were treated with 5 μM MG-132. For hyperosmotic treatment, cells were incubated in a medium supplemented with 0.9 mL growth medium and 0.1 mL 10× PBS immediately before imaging^71^. For hypoosmotic treatment, cells were incubated in a mixture of 0.5 mL medium and 0.5 mL MilliQ water immediately before imaging^71^. Imaging was done continuously for 20 min post-treatment, with solutions added directly on the microscope stage to predefined metaphase cell positions (details in Microscopy section).

### CENP-E reactivation

hTERT-RPE1 cells expressing CENP-A-GFP and centrin-GFP were incubated overnight with 20 nM SPY-DNA-650 (Spirochrome) and were acutely treated with 80 nM GSK-923295 (MedChemExpress, IC50 value 3.2 nM) 1 h before imaging. Just before imaging samples were washed three times with pre-warmed medium and were imaged continuously thereafter for 20 min with 10s interval (details in Microscopy section). Every third frame was analyzed. Cells with ∼10 polar chromosomes were chosen for imaging.

### Sample preparation for live cell imaging

Cells were seeded and cultured in DMEM with supplements at 37°C and 5% CO_2_ on uncoated 35-mm glass coverslip dishes with 0.17-mm (1.5 coverglass) glass thickness (MatTek Corporation). To visualize microtubules in U2OS and RPE1 cells expressing CENP-A-GFP and centrin1-GFP, SPY650-tubulin dye (*λ*_Ex_ 652 nm, *λ*_Em_ 674 nm) (Spirochrome AG) was introduced to the dishes at a final concentration of 100 nM, 2–3 hours before imaging. Additionally, for chromosome visualization in compression experiments and determination of mitotic phases, 50 μL of NucBlue Live Ready Probes Reagent (Hoechst 33342) dye (Thermo Fisher Scientific) was added to the dishes 1 minute before imaging.

### Live-cell microscopy

The U2OS and hTERT-RPE1 cell lines expressing CENP-A-GFP and centrin1-GFP were imaged for compression experiments were imaged using the Bruker Opterra Multipoint Scanning Confocal Microscope (Bruker Nano Surfaces). The system was mounted on a Nikon Ti-E inverted microscope equipped with a Nikon CFI Plan Apo VC ×100/1.4 numerical aperture oil objective (Nikon). During imaging, cells were maintained at 37°C in an Okolab Cage Incubator (Okolab). A 22-μm slit aperture was used, and the xy-pixel size was set to 83 nm. GFP and mCherry fluorescence were excited using 488-nm and 561-nm diode laser lines, respectively, while SPY dyes were excited using a 640-nm diode laser line. Opterra Dichroic and Barrier Filter Set 405/488/561/640 were employed to separate excitation light from emitted fluorescence. Images were captured with an Evolve 512 Delta EMCCD Camera (Photometrics) with no binning. To cover the entire metaphase spindle, z-stacks were acquired at 30–60 focal planes separated by 0.5 μm using unidirectional xyz scan mode. The system was controlled with Prairie View Imaging Software (Bruker Nano Surfaces).

hTERT-RPE1 parental cells and its post-tetraploid clones expressing H2B-GFP as well as human hTERT-RPE1 cells with or without p53KD and with expression of H2B-Dendra, RPE1 cells stably expressing CENP-A-GFP and centrin1-GFP treated with CENP-E inhibitor, and U2OS cells for MUG experiments, were imaged using the Lattice Lightsheet 7 system (Zeiss), as described previously^49^. The system was equipped with an illumination objective lens 13.3×/0.4 (at a 30° angle to coverglass) with a static phase element and a detection objective lens 44.83×/1.0 (at a 60° angle to coverglass) with an Alvarez manipulator. Automatic water immersion was applied from the motorised dispenser at intervals of 20 or 30 minutes. Immediately after sample mounting, the ‘create immersion’ option was applied four times, followed by ‘auto immersion’ at 25-minute intervals. The sample was illuminated with a 488 nm diode laser (power output 10 mW) with laser power set to 1–2%. Detection was performed using a Hamamatsu ORCA-Fusion sCMOS camera with an exposure time set to 15–20 ms. The LBF 405/488/561/642 emission filter was used. During imaging, cells were maintained at 37°C with 5% CO_2_ in a stage incubation chamber system (Zeiss). The imaging area width in the x dimension was set from 1 to 1.5 mm. The time interval between consecutive frames was set to 2 minutes and in CENP-E inhibitor washout experiments to 10 seconds.

Confocal live-cell imaging of RPE1 cells stably expressing CENP-A-GFP and centrin1-GFP, treated with nocodazole, hyperosmotic, hypoosmotic, or left untreated, was performed on a Dragonfly spinning disk confocal microscope (Andor Technology) using a 100×/1.47 HC PL APO oil objective (Nikon) and a Sona 4.2B-6 Back-illuminated sCMOS camera (Andor Technology), acquisition mode set to high speed. Images were acquired using Fusion software. During imaging, cells were maintained at 37 °C and 5% CO₂ in a heated chamber (Okolab). GFP and SPY-DNA-650 were excited using the 488-nm and 637-nm laser lines, respectively, with corresponding emission filters. Z-stacks of 21 focal planes were acquired at 1 µm intervals every 30 s at 4–6 predefined metaphase cell positions simultaneously.

### Sample preparation for immunofluorescence

For labelling centrioles and microtubules in RPE1 cells grown on glass-bottom dishes (14 mm, No. 1.5, MatTek Corporation), a pre-extraction step was carried out with pre-warmed PEM buffer at 37°C (0.1 M PIPES, 0.001 M MgCl_2_ × 6 H2O, 0.001 M EDTA, 0.5% Triton-X-100) for 30 seconds at room temperature, followed by fixation with 1 mL of ice-cold methanol for 2 minutes. After fixation, cells underwent three washes of 5 minutes each with 1 mL of PBS and were permeabilized with 0.5% Triton-X-100 solution in water for 30 minutes at room temperature. Subsequently, to block nonspecific binding, cells were incubated in 1 mL of blocking buffer (2% normal goat serum, NGS) for 2 hours at room temperature. After three more washes of 5 minutes each with 1 mL of PBS, cells were incubated with 500 μl of primary antibody solution overnight at 4°C. Antibody incubation was carried out using a blocking solution composed of 0.1% Triton and 1% NGS in PBS. The following primary antibodies were used: rabbit anti-centrin-3 (1:500, Abcam, ab228690) and rat anti-tubulin (1:500, MA1-80017, Invitrogen). Secondary antibodies used were donkey anti-rabbit Alexa Fluor 647 (1:1000, Abcam, ab150075) and donkey anti-rat Alexa Fluor 594 (1:1000, Abcam, ab150156). Finally, cells underwent three 10-minute washes with 1 mL of PBS before imaging, either immediately or after being stored at 4°C for a maximum period of one week. For cytokinesis failure experiments in RPE1 cells, 5 µM MG132 (Merck) for 1 hour prior to fixation.

For imaging of the mitotic spindle in HCT116-derived cells, the cells were grown on black glass-bottomed 96-well plates and treated with 10 µM MG132 (Calbiochem) for 3 hours prior to fixation. Cells were briefly washed with PBS, fixed with 100% methanol for 10 min at −20°C, washed three times with PBS, blocked with using 5% FCS/PBS solution with addition of 0.5% Triton-X-100, incubated overnight with anti-α-tubulin mouse (1:500, Sigma-Aldrich, CP06) and anti-γ-tubulin goat (1:1000, Santa Cruz, sc-7396-R), washed with 400 mM NaCl and subsequently twice in PBS-T, and detected with Goat anti-mouse DyLight 594- and DyLight 649-conjugated antibodies (Abcam, ab96873 and ab96886, respectively). DNA was stained with SYTOX Green with added RNase. Cells were finally washed in 400 mM NaCl and twice in PBS-T.

Organoids were fixed either in warm 4% paraformaldehyde or ice-cold methanol for 1 hour. Subsequent permeabilization, blocking, and staining procedures were performed as described previously^113^. Primary antibodies, rabbit anti-centrin-3 (1:300, Abcam, ab228690) and rat anti-tubulin (1:100, MA1-80017, Invitrogen), were diluted in the working buffer and incubated overnight at 4°C. Secondary antibodies used were: donkey anti-rabbit Alexa Fluor 488 or 594 (1:500, Abcam, ab150061 and ab150064, respectively) and donkey anti-rat Alexa Fluor 594 (1:500, Abcam, ab150156), which were also incubated overnight at 4°C. Staining with DAPI (1 µg/mL) and SiR-actin (1 µM or 2 µM, Spirochrome) was performed for 20 minutes at room temperature.

### Fixed-cell microscopy

All fixed RPE1 cells were imaged by the LSM 800 confocal system (Zeiss) with following parameters: xy sampling, 1.58 μm; z step size, 0.5 μm; total number of slices, 26; pinhole, 45– 56 μm; unidirectional scan speed, 10; averaging, 1; 63x Oil DIC f/ELYRA objective (1.4 NA); GaAsP detectors. Laser lines of 405 nm, 561 nm, and 640 nm were used for excitation. Image acquisition was performed using ZEN 3.5 software (blue edition; Zeiss). All fixed HCT116-derived cells were imaged by the Marianas SDC system with inverted Zeiss Observer.Z1 microscope. Following parameters were used: z-step size 0.5 μm, Plan Apochromat 63x magnification oil objective, with laser lines 473 nm, 561 nm and 660 nm (LaserStack, Intelligent Imaging Innovations, Inc., spinning disc head (Yokogawa), CoolSNAP-HQ2 and CoolSNAP-EZ CCD cameras (Photometrics, Intelligent Imaging Innovations, Inc.). Image acquisition was performed using SlideBook 5.1.

Confocal microscopy of fixed mSI organoids was conducted using the Expert Line easy3D STED microscope system (Abberior Instruments) with following parameters: 100x/1.4NA UPLSAPO100x oil objective (Olympus) and avalanche photodiode (APD) detector. Laser lines of 405 nm, 488 nm, 561 nm, or 647 nm were utilized for excitation. Image acquisition was performed using the in Imspector software. The xy pixel size was set at 100 nm and 200 nm, and z-stacks were obtained with 500 nm and 2 µm intervals for individual cells and for whole organoids, respectively.

### Measurements of spindle and metaphase plate parameters

In cell lines, spindle length was determined in ImageJ from maximum intensity projections (MIPs) or sum-intensity projections of z-stacks of individual cells, as the distance between spindle poles, which were identified as the center of the centriole pair or, if centrioles were absent, as the center of a prominent tubulin signal. Spindle width was defined as the distance between the outermost tubulin-labelled bundles in fixed cells where opposing centrosomes were within 7 µm in the z-dimension or as the distance between the outermost kinetochores in fixed CENP-E-depleted, live-imaged CENP-E-reactivated cells, control untreated, nocodazole-treated, and osmotically-shocked cells. Metaphase plate width and thickness were quantified on MIPs or sum-intensity projections of z-stacks of individual cells in both cell lines and tumor PDOs. Plate width was measured as the distance between the outermost chromosome ends perpendicular to the spindle axis and plate thickness along the spindle axis. These measurements were performed in cells with opposing centrosomes within 7 µm or 3 µm in the z-dimension, respectively. Cells with chromosomes in the region between the pole and the metaphase plate and those with multiple polar chromosomes were excluded from plate thickness measurements. Single polar chromosomes were not considered as part of the plate. In eukaryotes, spindle length and width, as well as metaphase plate width and thickness, were measured from published images (Extended data Table 1) as described above for cell lines.

In a cross-section of the metaphase plate, chromosome arms stick out, resulting in an irregular shape of the plate. We estimated the effective volume of the metaphase plate by approximating the shape of the plate with a cylinder. This approach allowed us to analyze published experimental data, many of which lack information about the three-dimensional shape of the spindle. Approximation of the plate shape with a cylinder was validated in previous studies on diverse systems including: mouse embryonic stem cells, Ptk2 cells^22^, human HeLa and RPE1^32,72,114^, and multiple Ichthyosporea species^115^. We estimated the metaphase plate volume in different eukaryotic species from the measured width (W) and thickness (T) using the formula for a cylinder: 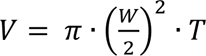. Metaphase plate density across species was calculated as the ratio between genome size and the estimated volume of the metaphase plate. Angles between poles, discriminated either by centrin signal or center of tubulin signal, and outermost kinetochore fibers were measured using *Angle tool* in ImageJ on MIPs. In freshly generated or naturally present polyploids the metaphase plate thickness and width were measured only in cells where bipolar spindles were formed, evidenced from the clustered centrosomes or a single metaphase plate, and rarely in PDOs when one dominant metaphase plate was formed in a multipolar spindle. The number of metaphase chromosomes in CENP-E depleted cells was determined by subtracting the number of chromosomes outside the metaphase plate from the total number of chromosomes in diploid human cells (2n = 46), based on kinetochore signals. Nuclear area was assessed in ImageJ using the polygon selection tool on MIPs or sum-intensity projections of z-stacks of individual cells in the DNA channel. For live-imaged cells, the nuclear area was measured a few hours before nuclear envelope breakdown (NEBD) in the DNA channel. Spindle height was measured in ImageJ as the distance between the first and last z-frame showing tubulin signal. This measurement was corrected by multiplying the obtained spindle height by a factor of 0.81 to account for refractive index mismatch^114^. In PDOs, we measured all cells that entered metaphase, excluding only those that were highly tilted.

Metaphase plate thickness was not quantified after CENP-E depletion in fixed cells, for the following reasons. CENP-E depletion activates the spindle assembly checkpoint and prolongs mitosis, causing increased microtubule pulling and chromosome dispersion within the metaphase plate^49^. Additionally, the extent of metaphase oscillations decreases as cells progress closer to anaphase^108^, producing effectively higher plate thickness in prometaphase cells. The longer spindle length in cells with more unaligned chromosomes likely reflects a prometaphase-like state^49^, consistent with previous reports that spindle length decreases from prometaphase to metaphase^69^.

In mouse primary cells, analyzed cells in metaphase comprised either single cells or portions of early organoid formations from the 2-cell stage onwards. Only multicellular structures lacking the characteristic asymmetric aspect ratios seen in mature organoids undergoing apical mitosis were considered. Selection criteria were based on nuclear and chromosome positioning relative to actin-labelled cell boundaries in images of complete organoid structures. Metaphase cells were categorized into two groups: diploid or polyploid, determined by the presence of either 2 or more centrin-2 or centrin-3 pairs, respectively. Spindle width, metaphase plate thickness, and diameter were quantified as described for cell lines. Spindle length was measured as the three-dimensional distance between opposing centrosomes or midpoint positions of centrosome clusters, as indicated by the centrin signal. Except for spindle length (p = 0.025), there were no significant differences observed in diploid or polyploid groups between cells fixed with methanol or paraformaldehyde, thus they are collectively presented. Across all parameters, no significant differences were noted within diploid or polyploid groups between cells derived from centrin-2-labelled or unlabeled organoids, or between cells originating from organoids treated or untreated with the ROCK inhibitor, hence they are represented together.

Metaphase plate width and spindle length in acentriolar cells were measured from published images using ImageJ, with the scale bar serving as the distance guide. Comparisons were limited to cells processed and imaged using the same protocols, including live-imaged human oocytes, mouse embryonic cells, and mouse and human oocytes fixed by the same methods, as referenced in Extended Data Table 2. For HAP1 cells, data were obtained from representative examples of naturally present near-haploid and near-diploid subpopulations, as detailed in the reference provided in Extended Data Table 2.

Cell area in cell lines was measured in ImageJ using the Polygon selection tool on maximum intensity projections of the tubulin channel, with the cytoplasmic tubulin boundary defining the cell perimeter. Measurements were manually verified by checking that the tubulin-based boundary corresponded to the centrin-based boundary in the other channel. Total chromatin intensity of metaphase plate in cell lines was calculated using the formula: Integrated density in metaphase plate – (Mean background intensity × Area of measurement). Background intensity was measured from three independent regions outside the cell, and the mean background was subtracted from the total DNA signal. Chromatin density was then calculated as the background-corrected total DNA intensity divided by the measured metaphase plate area. Chromatin density was measured using the same approach following CENP-E reactivation in live cells. In CENP-E-reactivated cells, the metaphase plate area was defined in every third frame by the continuous stretch of metaphase chromatin, and successful chromosome entry into the plate was defined as the frame in which both sister kinetochores became part of a continuous plate signal. For control and MUG U2OS cells, cell area was measured in ImageJ using the same approach in the tubulin channel in the last frame before anaphase.

Tubulin intensity for control and nocodazole-treated cells in the metaphase plate was measured on sum-intensity projections within the metaphase plate region defined by the DAPI signal. Background was subtracted based on the average of three measurements taken within the same cell, outside the spindle region. Metaphase plate and spindle dimensions were measured on MIP images in nocodazole and osmotic shock experiments. The analysis was performed on every frame in which all structures were visible, which resulted in a smaller number of analyzed frames due to sudden spindle collapse in nocodazole experiments.

### Image processing, data analysis and statistics

Image analysis was performed in ImageJ (National Institutes of Health) and plotting was done in Python (The Python Software Foundation) and MatLab (MathWorks). For plotting univariate scatter plots, Matlab “UnivarScatter” extension was used (https://github.com/manulera/UnivarScatter). Raw images were used for quantification. Figures were composed in Adobe Illustrator CS5 (Adobe Systems). In Extended Data Fig. 2g (first and middle bottom), diploid and hypotetraploid groups contain both DMSO- and centrinone-treated cells. In Extended Data Fig. 2g (middle top), all groups contain both 1:0 and 1:1 spindles, while Extended Data Fig. 2g (middle bottom) contains only 1:1 spindles, as the number of centrioles significantly influences only spindle length (see Extended Data Fig. 3a-c). In Fig. 2c and Extended Data Fig. 2g (bottom) and i, the tetraploid group contains both cells generated by cytokinesis failure and mitotic slippage whereas the diploid group contains both H2B-Dendra2 and H2B-GFP expressing RPE1 cells.

Bioinformatics analysis of gene expression data from diploid RPE1 and post-tetraploid RPT clones^38^ was performed in R/Bioconductor using “limma” for normalization and differential expression analysis. Log2 expression ratios (RPT compared to RPE1) were calculated for each gene. Processed RNA sequencing data were analyzed using Pathway Analysis, with “fdrtool” for false discovery rate correction and “ggplot2” for visualization. Genes were filtered using UniProt identifiers to focus on kinesin superfamily proteins (KIFs), spindle assembly factors, and MAPs, guided by GOCC terms.

The p values when comparing two independent classes were obtained using the independent two-sided Student’s t-test (significance level was 5%). The p values when comparing multiple classes were obtained using the one-way ANOVA test followed by Tukey’s Honest Significant Difference test (significance level was 5%). For testing the significance of the fits, the t-test of the slope coefficient in the linear regression was used (significance level was 5%). For testing the significance of relationships between variables, Pearson correlation coefficients were calculated and their p-values determined using the t-test for correlation. The p values and z-score when comparing the difference between two dependent correlations with one variable in common were obtained using Steiger’s correlation Z-test^116^ (significance level was 5%). No statistical methods were used to predetermine the sample size. The experiments were not randomized and, except where stated, the investigators were not blinded to allocation during experiments and outcome evaluation.

## Acknowledgements

We thank Alexey Khodjakov, Helder Maiato, and Rene Medema for providing cell lines; the Laboratory for Neurodegenerative Disease Research led by Silva Katušić at the Ruđer Bošković Institute for assistance with isolation of mouse small intestine; Ivana Šarić for the illustrations and assembling the figures; and members of the Tolić and Pavin groups for their constructive comments on the manuscript. This work was funded by the European Research Council (ERC-SyG 855158, I.M.T., N.P., and G.J.P.L.K.); the Croatian Science Foundation projects (HRZZ IP-2019-04-5967, N.P.; HRZZ IP-2022-10-1091, M.LJ. and J.T.; HRZZ IP-2024-05-5336, I.M.T.) and Swiss-Croatian Bilateral Project (IPCH-2022-10-9344, I.M.T.); projects co-financed by the Croatian Government and the European Union through the European Regional Development Fund under the Competitiveness and Cohesion Programmes 2014-2020 and 2021-2027, via the IPSted project (KK.01.1.1.04.0057, I.M.T.), and the project ‘Implementation of cutting-edge research and its application as part of the Scientific Center of Excellence for Quantum and Complex Systems, and Representations of Lie Algebras’ (PK.1.1.10.0004, N.P., J.T., M.N., M.LJ., I.M.T.), and the Deutsche Krebshilfe (Project 70115224, Z.S.). This research was performed using services, storage, and computing resources provided by the University of Zagreb University Computing Center - SRCE.

## Contributions

Analysis of spindle scaling across eukaryotes was conceptualized and carried out by I.M.T. and N.P., with help from K.V. and L.G. Experiments on RPE1 cells across ploidies were performed and analyzed by L.G., K.V., and A.P., with help in analysis from L.P. Transcriptome analysis and HCT116 movies were done by A-L.H. and Z.S. Patient-derived organoid movies from G.J.P.L.K.’s lab were prepared by T.V.R. and analyzed by K.V. Primary mouse cell experiments were carried out by I.D. on organoids established by Ma.T. CENP-E depletion experiments were conducted by K.V. and analyzed by L.P. Compression and MUG experiments were performed by Mo.T. and L.G. Nocodazole, tonic shock and CENP-E reactivation experiments were performed and analyzed by K.V. and L.G. The theoretical model was designed by N.P. with help from M.N., J.T., and M.LJ., and solved by J.T., M.LJ., and M.N. The manuscript was written by I.M.T. and N.P. with help from K.V., L.G., and M.N., and feedback from all authors.

## Data availability

All relevant data supporting the findings of this study are provided within the article and its Extended Data files. Raw images used in this work are available from the corresponding authors upon reasonable request.

## Code availability

The code for the numerical computations of the theoretical model is available at https://github.com/matkec0/spindleFEM.

---

**Box 1.**
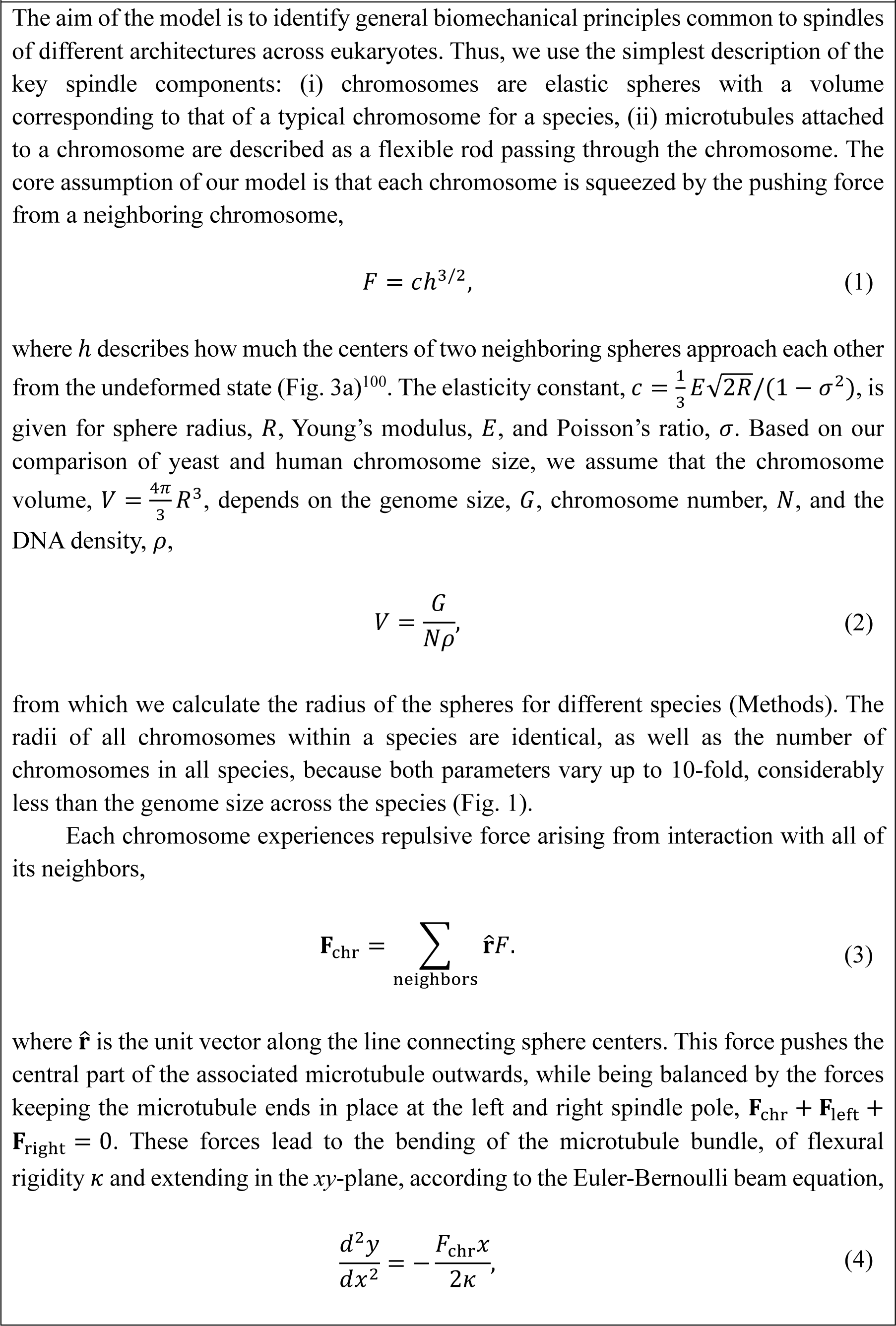
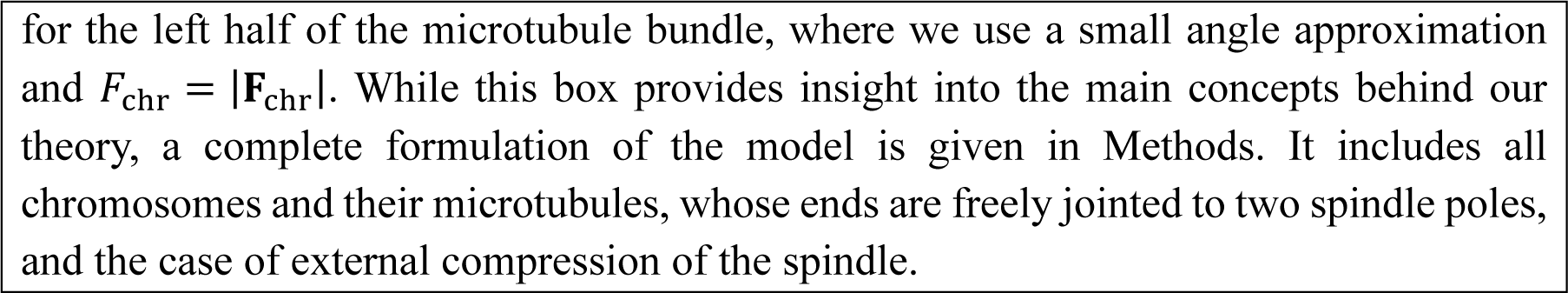
Model for spindle scaling.

## Extended Data

**Extended Data Fig. 1.**
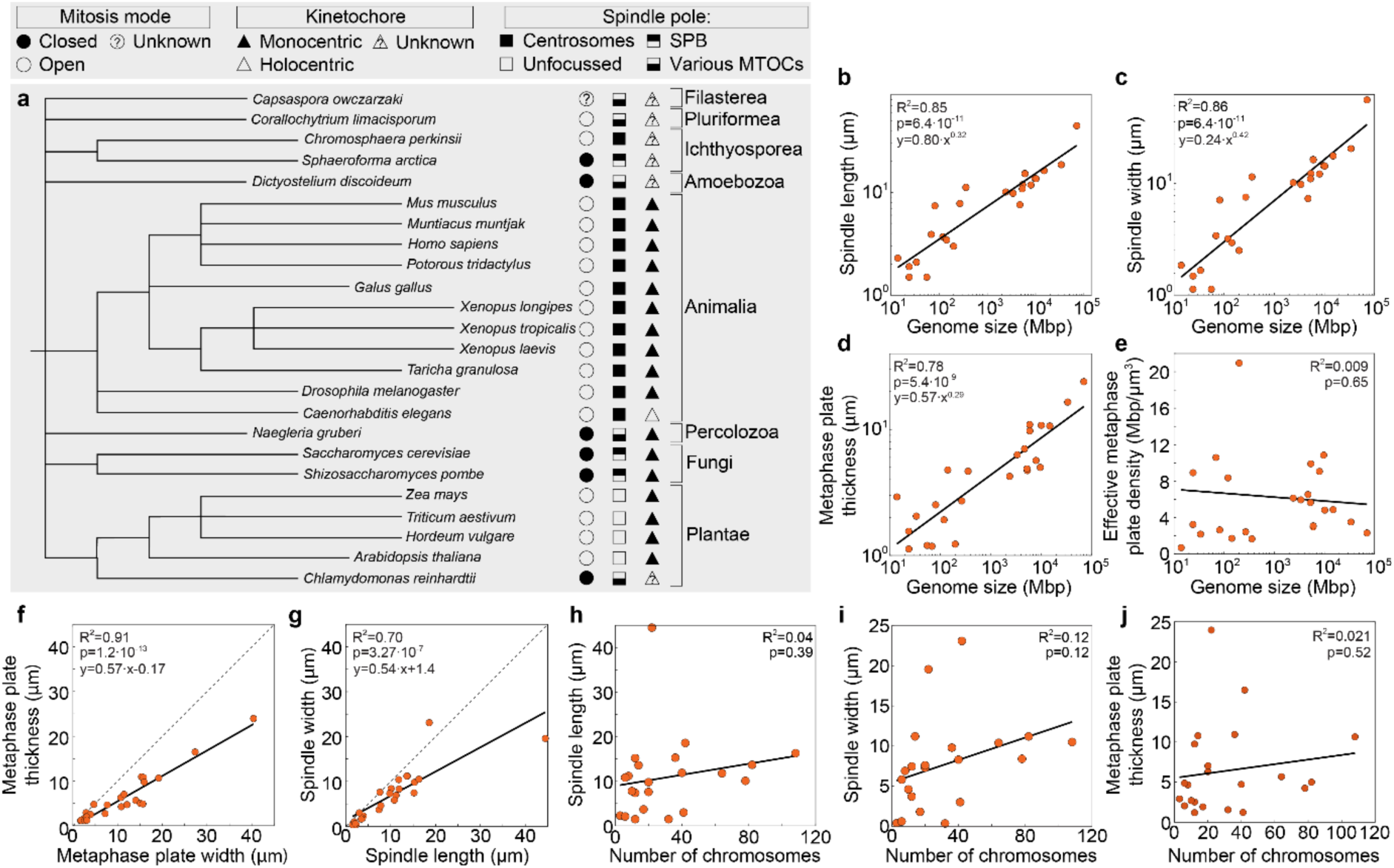
Phylogenetic tree of eukaryotic species used in the study and additional characterization of spindle and metaphase plate dimensions in relation to genome size and number of chromosomes. **a**, A phylogenetic tree based on the NCBI taxonomy, including species from Extended Data Table 1, with eukaryotic kingdoms, clades, or phyla denoted on the right. Schemes representing various modes of mitosis (circles), spindle pole organization (squares), and kinetochore structure (triangles) are shown according to the legend at the top. **b-d**, Spindle length (**b**), spindle width (**c**), metaphase plate thickness (**d**), as a function of the genome size, for the same species as in Fig. 1d, and a power-law fits. **e**, Effective metaphase plate density, calculated as genome size per volume of the metaphase plate, versus genome size, for the same species as in Fig. 1d, and a linear fit. **f**, **g**, Metaphase plate thickness versus metaphase plate width (**f**) and spindle width versus spindle length (**g**) for the same species as in Fig. 1d, and linear fits. Dashed lines represent y = x functions. **h**-**j** Spindle length (**h**), spindle width (**i**), and metaphase plate thickness (**j**) as a function of the chromosome number for the same species as in Fig. 1d, and a linear fits. Abbreviations: SPB, spindle pole body, MTOCs, microtubule-organizing centers. The p-values are obtained from a two-sided t-test on the regression slope.

**Extended Data Fig. 2.**
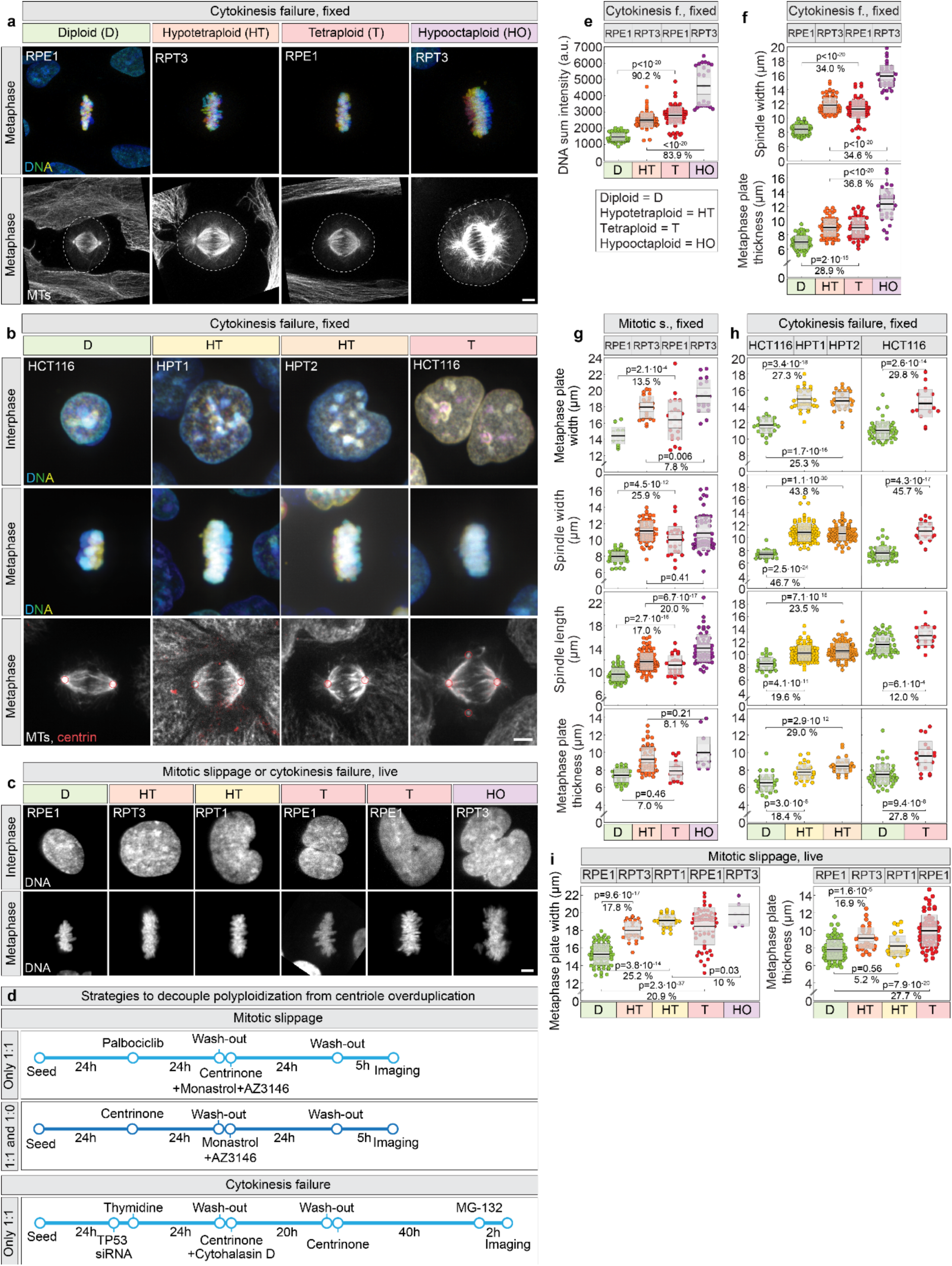
The scaling of spindle dimensions with ploidy in human differentiated cells is determined by the amount of DNA in the metaphase plate. **a**, DNA stained with DAPI (color-coded for depth) (top) and microtubules (MTs) immunostained with α-tubulin (gray) with circled cell bounds (bottom) in fixed RPE1 and RPT3 cells of different ploidy. **b**, DNA stained with DAPI (color-coded for depth) in fixed HCT116, HPT1 and HPT2 cells of different ploidies in interphase (top), metaphase (middle), and in metaphase with microtubules (MTs) immunostained with α-tubulin (MTs, gray) and centrioles with centrin-3 (red, encircled) (bottom). **c**, DNA in live-imaged RPE1, RPT1 and RPT3 cells of different ploidies stably expressing H2B-Dendra2 or H2B-GFP at interphase (top) and last frame before anaphase (bottom). **d**, Protocols to generate polyploid cells without centriole overduplication, targeting three outcomes: (i) mitotic slippage–induced polyploidization, yielding only cells with one centriole per centrosome (1:1) (top); (ii) mitotic slippage–induced polyploidization, yielding a mixture of cells with one centriole per centrosome (1:1) and cells with a single centriole (1:0) (middle); and (iii) cytokinesis failure–induced polyploidization, yielding only cells with one centriole per centrosome (1:1) (bottom). Exact concentrations are detailed in the Methods. **e**, DNA sum intensity in the metaphase plate of fixed metaphase cells of different ploidies after cytokinesis failure. **f**, Spindle width (top) and metaphase plate thickness (bottom) across fixed metaphase cells of different ploidies after cytokinesis failure. **g**, Metaphase plate width (top), spindle width (second), spindle length (third), and metaphase plate thickness (bottom) across fixed RPE1 and RPT3 cells of different ploidies produced by mitotic slippage. **h**, Metaphase plate width (top), spindle width (second), spindle length (third), and metaphase plate thickness (bottom) across fixed hypotetraploid HPT1 and HPT2 clones and their diploid HCT116 controls (left), and HCT116 tetraploids newly formed by cytokinesis failure, and their internal diploid controls (right). **i**, Metaphase plate width (left), and metaphase plate thickness (right) across live-imaged RPE1, RPT1 and RPT3 cells of different ploidies. Percentages on (**e**), (**f**), (**g**), (**h**) and (**i**) represent differences from the mean values of indicated groups; the thick black line shows the mean; the light and dark gray areas mark 95% confidence interval on the mean and standard deviation, respectively; p values from Tukey’s Honest Significant Difference test are given in (**e**), (**f**), (**g**), (**h**, left) and (**i**); p values from Student’s t-test are given in (**h**, right). Number of cells: (**e**) 86, 76, 80, 24; (**f**) 86, 76, 82, 42; (**g**) 15, 25, 24, 19 (top), 65, 113, 28, 60 (second, third), 37, 55, 20, 14 (bottom); (**h**) 32, 32, 32, 54, 20 (top), 32, 96, 100, 54, 20 (second), 32, 96, 100, 54, 20 (third), 32, 32, 32, 54, 20 (bottom); (**i**) 82, 36, 28, 63, 7 (left), 82, 36, 28, 63 (right); each group is a pool of at least three independent experiments. Abbreviations: a.u., arbitrary units; f., failure; s., failure. All scale bars, 5 µm.

**Extended Data Fig. 3.**
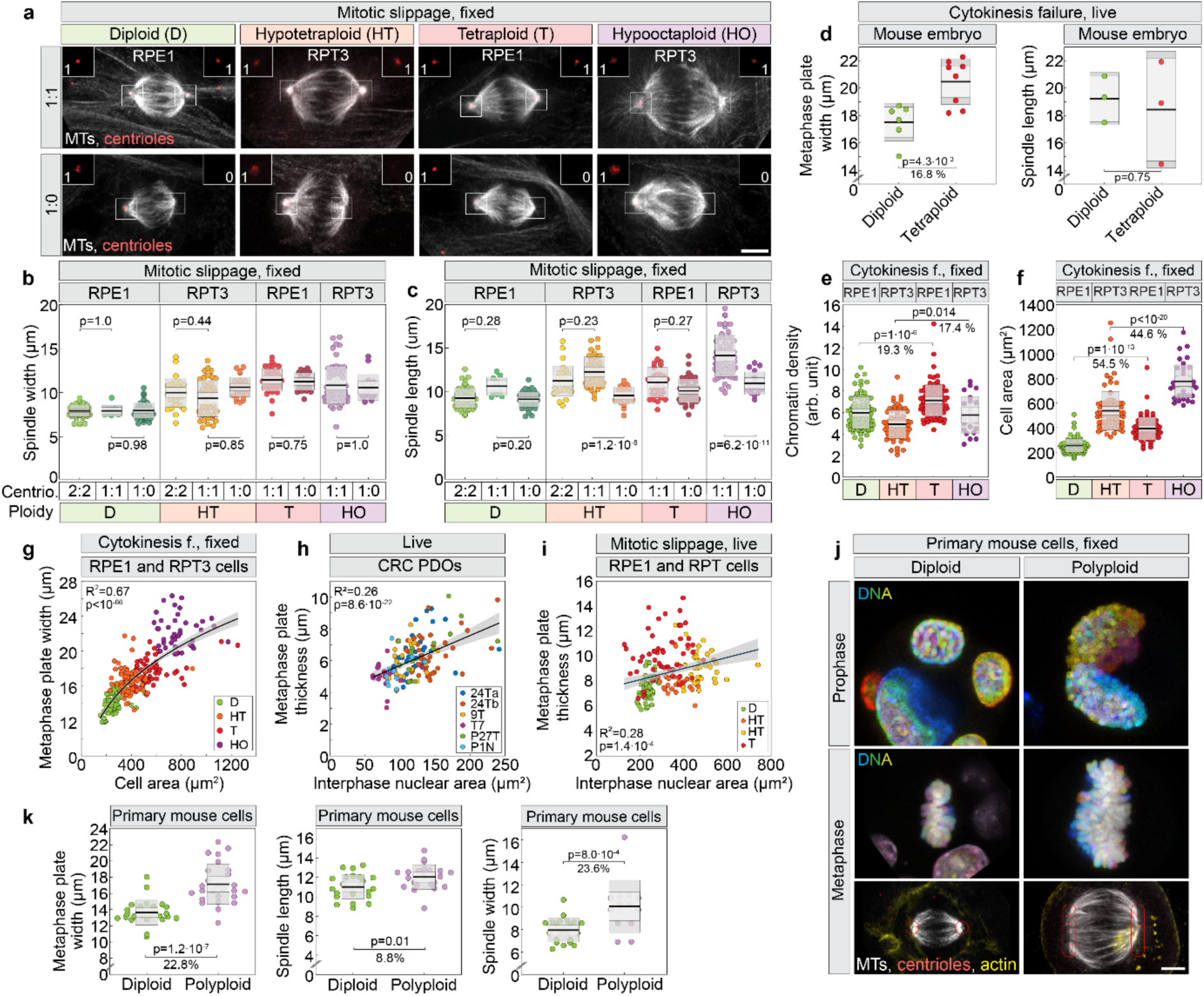
Metaphase plate width increases with increasing ploidy in human differentiated cell lines, tumor patient derived organoids, primary mouse cells and mouse embryonic cells, independently of centrioles. **a**, RPE1 and RPT3 cells of different ploidies immunostained for α-tubulin (MTs, gray) and centrin-3 (centrioles, red, encircled). The first row represents spindles with one centriole on each centrosome (1:1) and the bottom row represents spindles with only one centriole (1:0). Zoomed views of the centrosome area (boxed) are located in the upper corners. **b** and **c**, Spindle width (**b**), and spindle length (**c**) across different ploidies of RPE1 and RPT3 cells and different number of centrioles. The black line shows the mean; the light and dark gray areas mark 95% confidence intervals on the mean and standard deviation, respectively. **d**, Metaphase plate width (left), and spindle length (right) for diploid and tetraploid mouse acentriolar blastocysts. Data is measured from published images^43^. **e** and **f**, Estimated chromatin density in metaphase plate, measured as total fluorescence intensity per plate area (**e**), and metaphase cell area (**f**) across RPE1 and RPT3 cells of different ploidies. **g**, Metaphase plate width versus cell area in cells of different ploidies, with power-law fit and 95% confidence interval. **h**, Metaphase plate thickness versus interphase nuclear area of cells from live-imaged colorectal cancer (CRC) patient derived organoids (PDOs) across multiple lines (legend is defined in Fig. 2d caption). Black line represents mean linear fit; shaded area, 95% confidence interval. Metaphase plate thickness was measured at the last frame before anaphase onset. **i**, Metaphase plate thickness versus interphase nuclear area for live-imaged RPE1, RPT1 and RPT3 cells of different ploidies for conditions indicated in the legend. Orange dots represent RPT3 and yellow RPT1 cells. Black line represents mean linear fit; shaded area, 95% confidence interval. Metaphase plate thickness for live-imaged RPE1 cells was measured at the last frame before anaphase onset. **j**, DNA in primary mouse cells of different ploidy labelled with DAPI during prophase (top) and metaphase (middle), with immunostaining for α-tubulin (MTs, gray), centrin-3 (centrioles, red, encircled) and SiR-actin labelling (actin, yellow) (bottom). **k**, Metaphase plate width (left), spindle length (middle), and spindle width (right) in primary mouse cells of different ploidy. Number of cells: (**b**) 55, 9, 36, 27, 38, 21, 28, 40, 60, 16; (**c**) 55, 10, 41, 27, 38, 22, 28, 43, 63, 16; (**d** left) 6, 8; (**d** right) 3, 3; (**e**) 86, 76, 80, 24; (**f**, **g**) 86, 76, 79, 4; (**h**) 35, 44, 17, 20, 14, 10; (**i**) 58, 28, 36, 51; (**k** left) 26, 26; (**k** middle) 24, 22; (**k** right) 22, 13, each group is a pool of at least three independent experiments. In (**h**) and (**i**), lines represent mean linear fit; shaded areas, 95% confidence interval. P values from Tukey’s Honest Significant Difference test are given in (**b**), (**c**), (**e**) and (**f**); p values from Wald test are given in (**h**) and (**i**); p values from Student’s T-test are given in (**d**), (**g**) and (**k**). Abbreviations: centrio., centrioles; arb. Arbitrary; f. failure. All scale bars, 5 µm.

**Extended Data Fig. 4.**
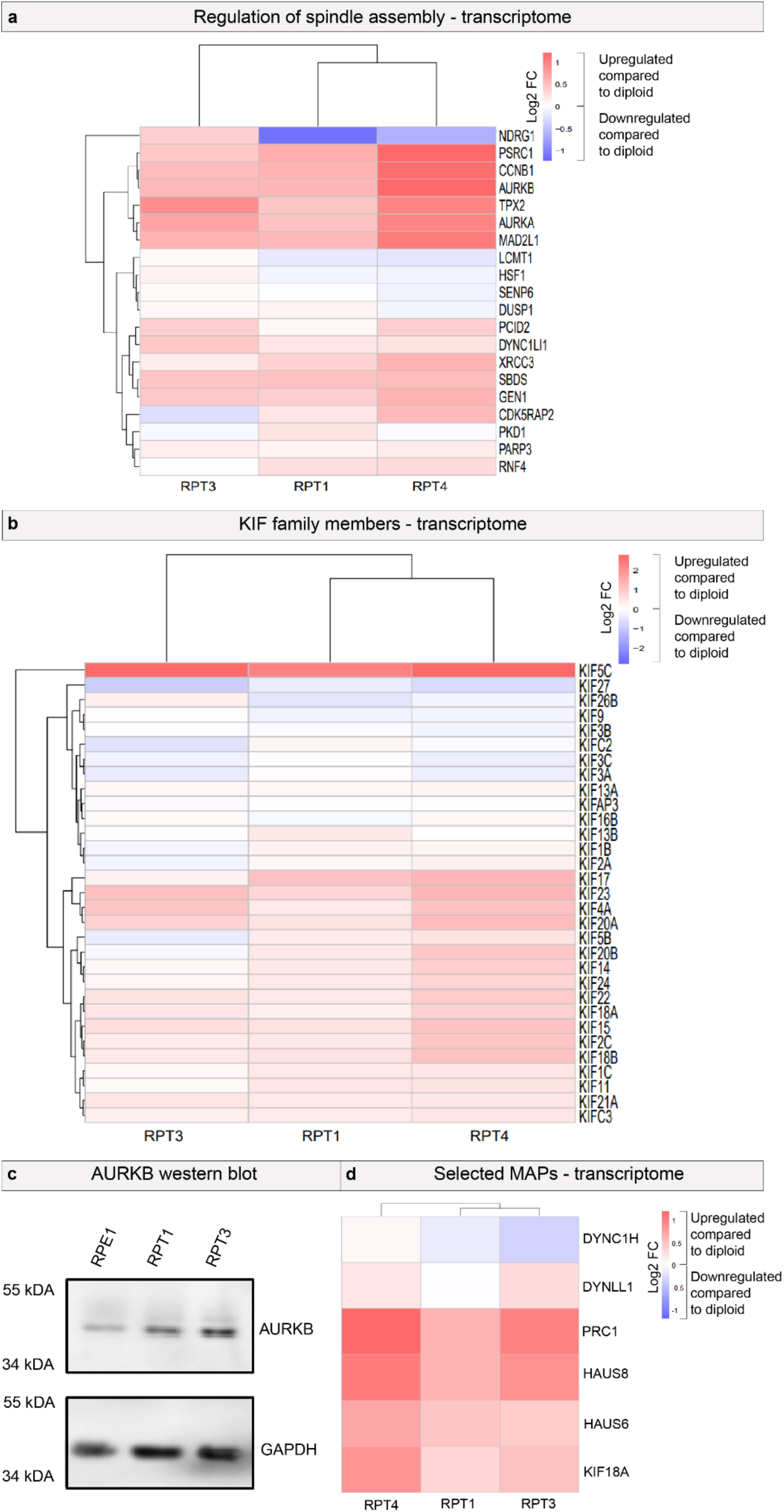
Transcriptome analysis reveals a sublinear increase of mitotic factors in adapted tetraploid RPE1 clones. **a**, Comparison of expression of genes involved in spindle assembly in hypotetraploid RPT cell lines normalized to their diploid parental wild type RPE1 counterparts. Genes significantly upregulated (red) and downregulated (blue) are shown (FDR < 0.05). FC, fold change. Note that logFC of ∼1 indicates an expression that matches ploidy status. **b**, Expression changes in genes of the kinesin (KIF) family from post-tetraploid RPT cell lines compared to the diploid parental wild type. GOCC annotation highlights the functional relevance of these changes, with significantly upregulated (red) and downregulated (blue) genes indicated (FDR < 0.05). GOCC annotation was employed to identify functional pathways. **c**, Immunoblot analysis of lysates from indicated cell lines. Antibody against Aurora B (AURKB) was used to estimate protein level, with glyceraldehyde-3-phosphate dehydrogenase (GAPDH) serving as the loading control. **d**, Comparison of genes coding for proteins which were perturbed on Fig. 4j in hypotetraploid RPT cell lines and their diploid parental wild type counterparts.

**Extended Data Fig. 5.**
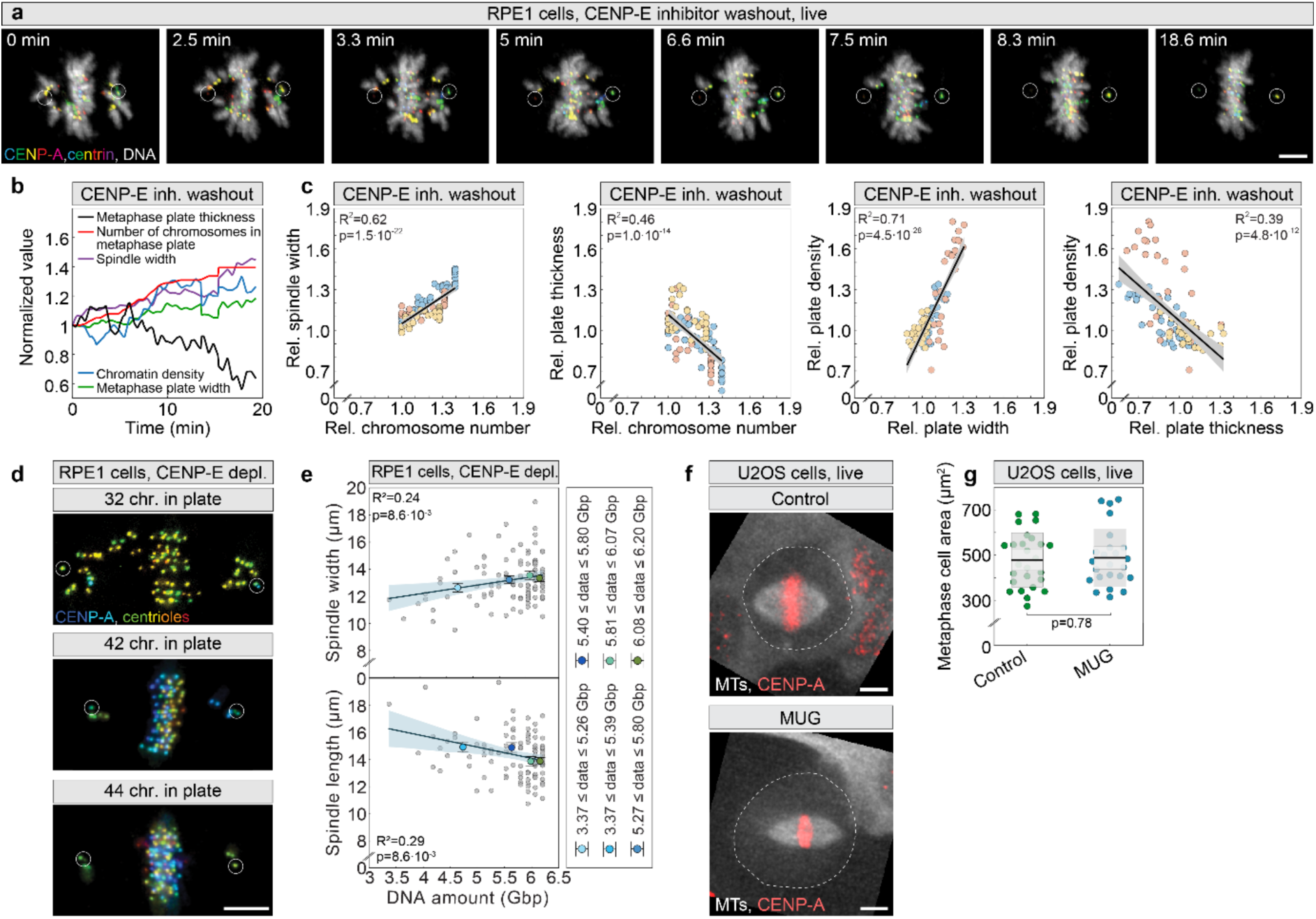
Decreased number of chromosomes in metaphase plate leads to narrower spindles. **a**, Timelapse of live-imaged RPE1 cell expressing CENP-A-GFP and centrin-3-GFP (color coded for depth with centrioles circled), and stained with SPY650-DNA (gray), after CENP-E inhibitor washout. Time after the washout is given. **b**, Relative mean curves for metaphase plate thickness and width, number of chromosomes in metaphase plate and chromatin density, in time after CENP-E inhibitor washout. Values were calculated relative to the initial value at the start of imaging. **c**, Relative spindle width (left), relative plate thickness (middle left) as a function of relative chromosome number after CENP-E inhibitor washout, metaphase plate width as a function of relative plate width (middle right), and relative plate density as a function of relative plate thickness (right) after CENP-E inhibitor washout. Values were calculated relative to the initial value at the start of imaging. Values from individual cells are colored differently. Black line, linear fit, gray area, 95% confidence interval. **d**, RPE1 cells stably expressing CENP-A-GFP, and centrin-3-GFP (centrioles, encircled, color coded for depth), after depletion of CENP-E, with indicated number of chromosomes in the metaphase plate. **e**, Spindle width (top) and spindle length (bottom) versus DNA amount in metaphase plate for RPE1 cells fixed after CENP-E depletion with mean ± SEM for conditions as indicated in the legend. The DNA amount was determined by multiplying the total genomic DNA in diploid human cells by the number of chromosomes retained in the metaphase plate. Black line, linear fit; blue area, 95% confidence interval. **f**, Spindles in live-imaged U2OS cells expressing CENP-A-GFP (red) and α-tubulin-mCherry (MTs, gray) in control (top) and mitosis with an unreplicated genome (MUG, bottom) cells. Cell bounds are denoted by a dashed line. **g**, Metaphase cell area for control and MUG cells. In (**g**), lines represent mean linear fit; shaded areas, 95% confidence interval. P values from Student t-test are given in (**g**) and (**c**) and from Wald test in (**e**). Number of cells: (**b**), (**c**) 3 cells from 3 independent experiments; (**e**) 115 (top), 105 (bottom) cells pooled from 2 independent experiments; (**g**) 26 and 25 cells, pooled from 2 independent experiments. Abbreviations: rel., relative; inh., inhibitor; rel., relative; depl., depletion; chr., chromosomes. All scale bars, 5 µm.

**Extended Data Fig. 6.**
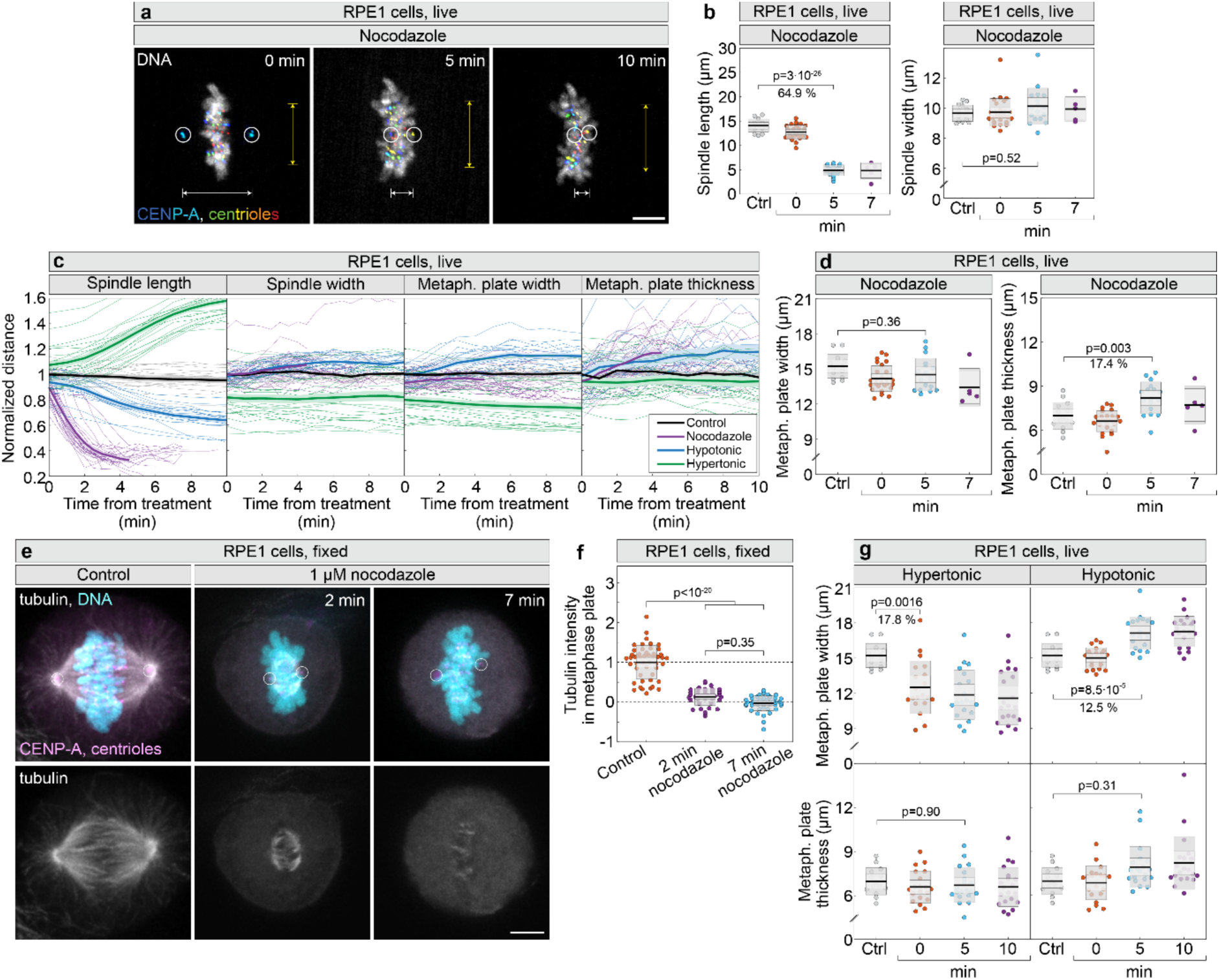
Spindle length is maintained by microtubules, in contrast to spindle width. **a**, Live-imaged RPE1 cells expressing CENP-A-GFP and centrin-3-GFP (color coded for depth with circled centrioles) and stained with SPY650-DNA (cyan) treated with 1 µM nocodazole. Times denote the start of imaging. Yellow arrows represent spindle width and white arrows spindle length. **b**, **d**, Spindle length (left) and spindle width (right) (**b**), and metaphase plate width (left) and metaphase plate thickness (right) (**d**), in analyzed frames, in control (Ctrl) and nocodazole-treated RPE1 cells. Percentages indicate differences between mean values of groups. Colored points represent individual cells; black lines show the mean, with light and dark gray areas marking 95% confidence intervals for the mean and standard deviation, respectively. **c**, Normalized values of spindle length (left), spindle width (middle left), metaphase plate width (middle right), and metaphase plate thickness (right) over time from treatment in RPE1 cells. Thick lines represent means, shaded areas represent SEM and small lines represent individual cells, colored per different treatments, as per legend on the right graph. **e**, Images of fixed RPE1 cells expressing CENP-A-GFP and centrin-GFP (magenta), immunostained for a-tubulin (gray), and labelled with DAPI (cyan). Time represents treatment duration. Only the tubulin channel is shown on the bottom. **f**, Tubulin intensity in the metaphase plate normalized to the mean of the control untreated group. Dispersion measures as in (**b**). **g**, Metaphase plate width (top) and metaphase plate thickness (bottom), in analyzed frames, in control (Ctrl) and hypertonically-treated (left) and control (Ctrl) and hypotonically-treated (right) RPE1 cells. Percentages indicate differences between mean values of groups. Dispersion measures as in (**b**). Number of cells: (**b** and **c**)16 and 29 in total per treatment; (**d**) 16, 29, 27, and 20 in total per treatment; (**f**) 48, 39, 45; (**g**) 27 and 20 cells in total per treatment, each group is a pool of at least three independent experiments. P values from Tukey’s Honest Significant Difference test are given in (**b**), (**c**), (**f**) and (**g**). Abbreviations: metaph., metaphase; Ctrl, control. All scale bars, 5 µm.

**Extended Data Fig. 7.**
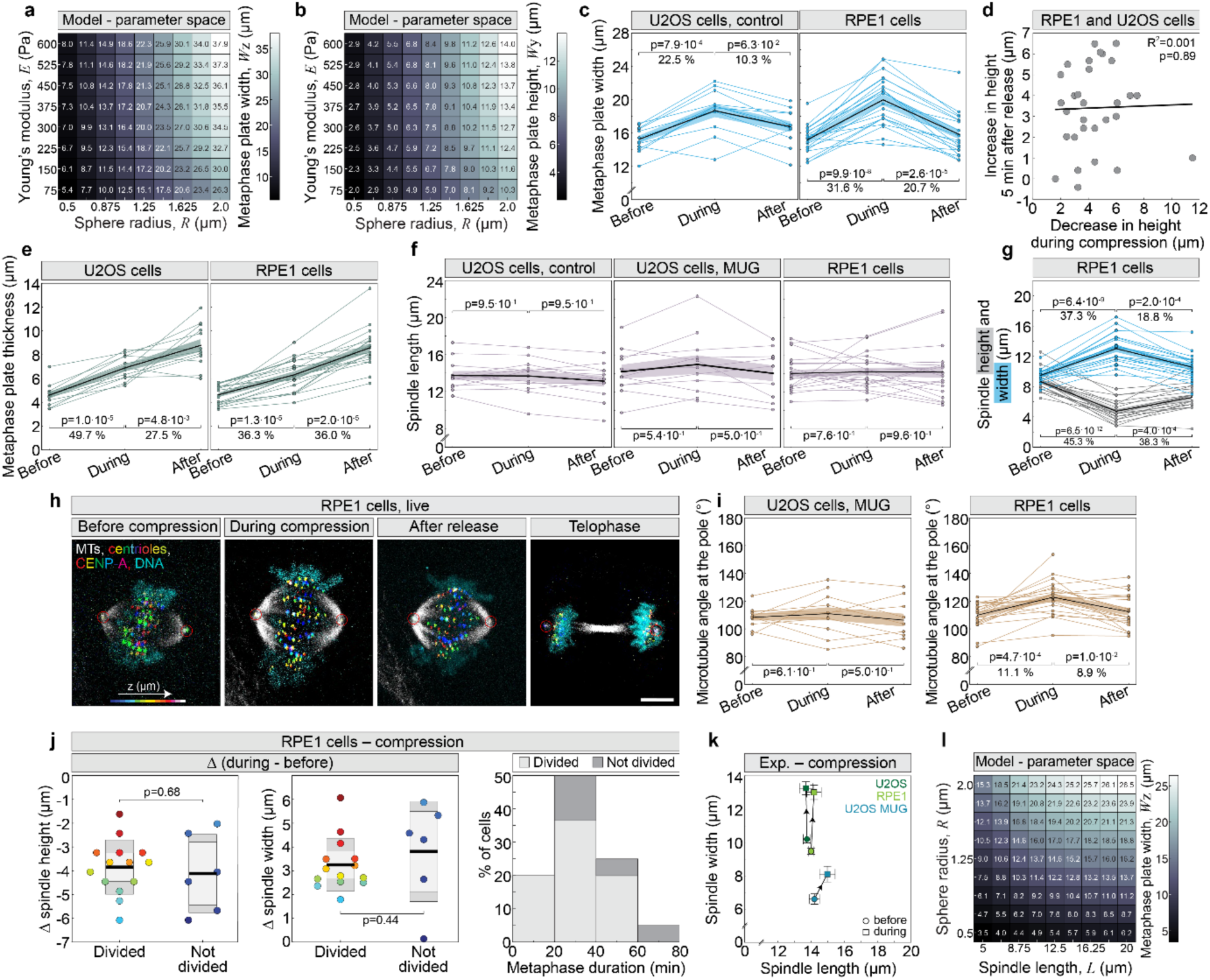
Acute spindle compression increases metaphase plate width and thickness without affecting spindle length and blocking cell proliferation. **a** and **b**, Phase diagram of the metaphase plate width, dimension perpendicular to the direction of external force, Wz (**a**), and the metaphase plate height, dimension in the direction of external force, Wy (**b**), as a function of Young’s modulus and sphere radius for compression coefficient *k* = 100 pN/μm. **c**, Metaphase plate width in U2OS control (left) and RPE1 (right) cells for three time points related to cell compression. **d**, Increase in spindle height 5 min after compression release, as a function of decrease in spindle height during compression, in both RPE1 and U2OS cells. Black line represents linear fit. **e**, Metaphase plate thickness for three time points related to compression of U2OS control (left) and RPE1 control (right) cells. **f**, Spindle length for three time points related to compression of U2OS control (left), U2OS mitosis with an unreplicated genome (MUG) (middle) and RPE1 control (right) cells. **g**, Spindle height (gray) and spindle width (blue) for three time points related to compression of control RPE1 cells. **h**, Time-lapse of control RPE1 cells stably expressing CENP-A-GFP and centrin-GFP (centrioles, encircled) (color-coded for depth) in compression experiment, with SiR-tubulin (MTs, gray) and Hoechst 33342 labelled DNA (cyan). **i**, Angle between pole and outermost kinetochore fiber for three time points related to compression of U2OS MUG (left) and control RPE1 (right) cells. **j**, Changes (Δ) in height (left) and width (middle) during compression, and metaphase duration histogram after compression for control RPE1 cells that either divided (light gray) or did not divide (dark gray) during imaging (right). Colored points on the left and middle represent individual cells; the black line shows the mean, with light and dark gray areas marking 95% confidence intervals for the mean and standard deviation. p-values from Student’s t-test are shown above. **k**, Mean ± SEM for spindle width versus spindle length for compression experiments; lines connect corresponding mean values before and after compression. Number of cells: 12, 11, 21. **l**, Phase diagram of metaphase plate width, *W*_*z*_, as a function of chromosome radius and spindle length, for *k* = 100 pN/μm. In (**c**), (**e**), (**f**), (**g**), and (**i**) black line, mean value; shaded area, 95% confidence interval. Number of cells - (**c**) 12 (left); 21 (right); (**d**) 28 in total; (**e**) 12 (left); 21 (right); (**f**) 12 (left), 21 (middle), 11 (right); (**g**) 21; (**i**) 21 (left), 11 (right); (**j**) 14, 6 (left); 14, 6 (middle); 20 (right); (**k**) 12 (left), 21 (middle), 11 (right). In (**a**), (**b**), and (**l**) parameters are as in Fig. 3, unless stated otherwise. Abbreviations: Exp., experiment. The number of cells equals the number of independent experiments.

**Extended Data Fig. 8.**
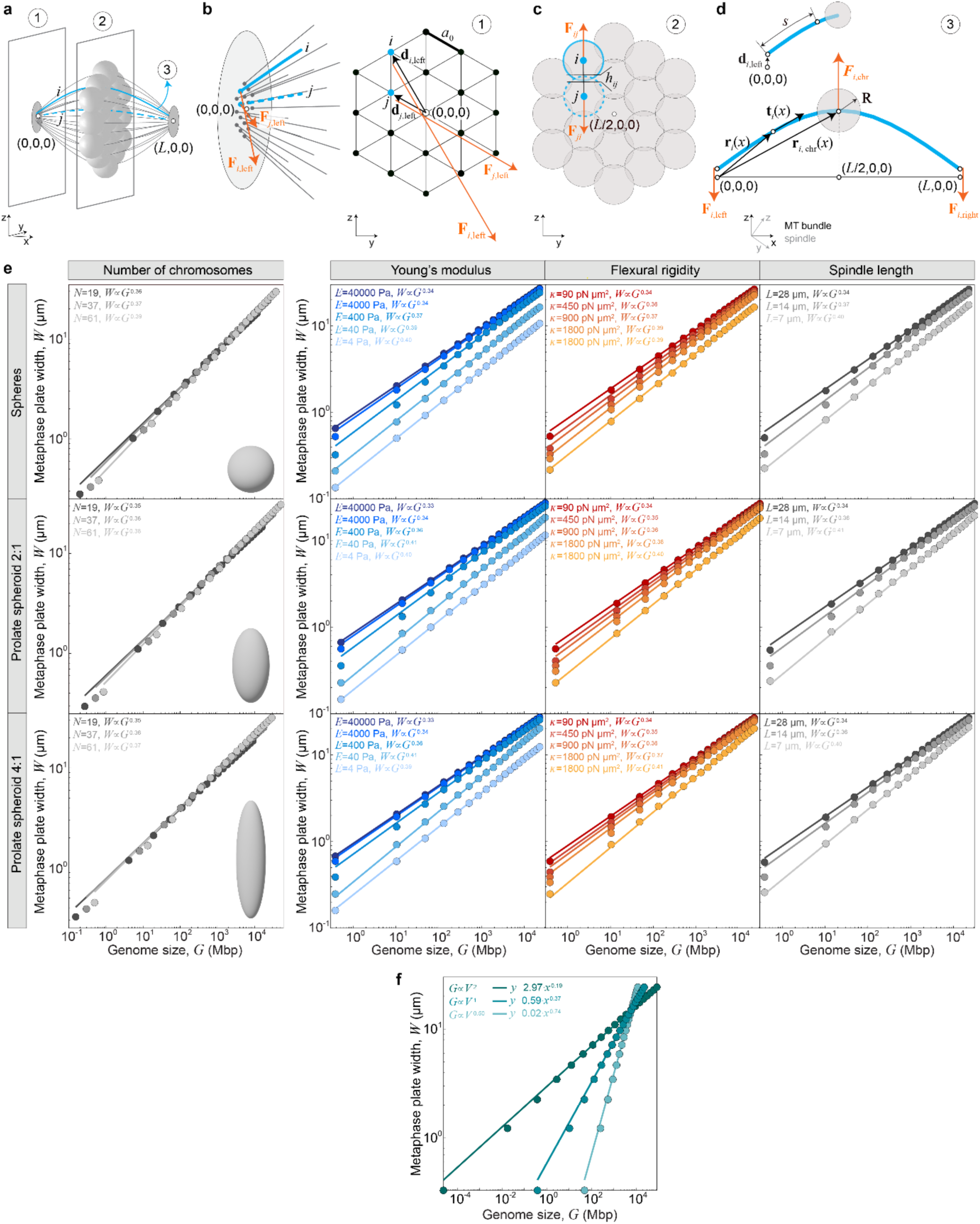
Scheme of the model. **a**, Three-dimensional scheme of the metaphase spindle with *N* = 19 chromosomes for the parameter values from Fig. 3. Two cross-sections of the spindle which contain the left spindle pole, and the metaphase plate are numbered (1) and (2) respectively, whereas a single bundle is numbered (3). Chromosomes are shown in light gray, microtubule bundles in gray, and spindle poles in dark gray. Two individual bundles, labeled *i* and *j*, are shown by solid and dashed blue lines, respectively. Coordinates of the right and left spindle poles are shown by triples of numbers. **b**, Scheme of the left spindle pole in side view (left) and end-on view (right). The forces exerted by the left pole on the microtubule bundles, *F*_*i*,left_ and *F*_j,left_, are shown in orange for bundles *i* and *j*, respectively. The positions of the left microtubule bundle ends are shown as blue dots for the bundles *i* and *j*, whereas other bundles are shown in black. The radius vectors for the left microtubule ends, *d*_*i*_ and *d*_j_, are shown in black, as well as the lattice parameter, *α_0_*. The coordinate of the left spindle pole is shown by a triple of numbers. **c**, Scheme of the metaphase plate in end-on view. The repulsive forces *F*_*i*j_ and *F*_j*i*_, mutually exerted between the bundles *i* and *j*, are shown in orange. The reduction of distance, *ℎ*_*i*j_ is shown in black. The coordinate of the metaphase plate center is shown by a triple of numbers. **d**, Scheme of the microtubule bundle *i*. The forces exerted by the pole on the left and right ends of microtubule bundles, *F*_*i*,left_ and *F*_*i*,right_ respectively, as well as the repulsive force exerted by the neighboring chromosomes, *F*_*i*,chr_ are shown in orange. The radius vector at the coordinate *x*, r_*i*_(*x*), the tangent, *t*_*i*_(*x*), the radius vector of the chromosome, r_*i*,chr_(*x*), and the radius of the chromosome, *R*, are shown in black. The coordinates of the left and right poles, as well as the center of the metaphase plate, are shown by triples of numbers. The contour length for bundle *i* is shown in the inset. The coordinate systems of the entire spindle and its individual components are shown in insets within each panel. **e**, Metaphase plate width, *W*, as a function of genome size, *G*, for varying shape of prolate spheroids, chromosome number, *N*, Young’s modulus, *E*, flexural rigidity, *κ*, and spindle length, *L*, shown on a log-log scale. The shape of prolate spheroids is parametrized by the ratio of equatorial to polar radius, *R_2_*/*R_1_* and it is depicted inside the left panel as a gray spheroid. The polar radius is held constant at *R_1_* = 1.25 μm. Rows (top to bottom) correspond to spheroid shapes with *R_2_*/*R_1_* = 1: 1, 2: 1 and 4: 1. Columns (left to right) show the variation of parameters *N*, *E*, *κ*, and *L*, respectively. **f**, Metaphase plate width, *W*, as a function of genome size, *G*, shown on log-log scale, assuming a nonlinear relationship between genome size and chromosome volume given by *G* = *G_0_*(*V*/*V_0_*)^*α*^. The exponent is varied among *α* = 0.5, 1 and 2, whereas the parameters *G_0_* and *V_0_* are fixed to the human genome size and volume of a sphere with *R* = 1.25 μm. In all panels parameters are as in Fig. 3, except the value of lettice parameter, *α*_0_ = 0.0025 μm.

## Extended Data Tables 1 and 2

**Extended Data Table 1.**
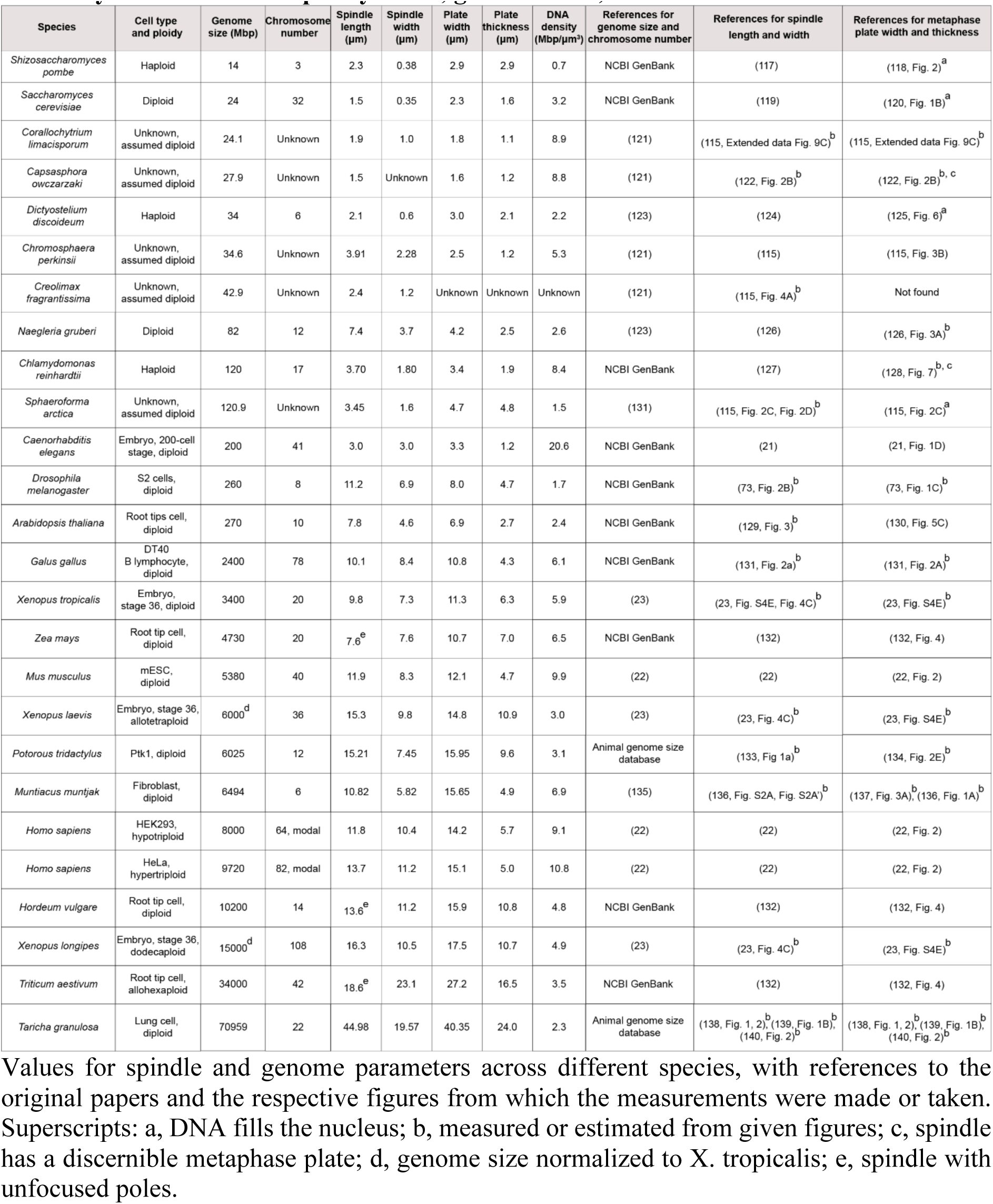
Measurements of metaphase plate width and thickness, spindle length and width, and metaphase plate density from published images of mitotic spindles in eukaryotes with diverse ploidy levels, genome sizes, and chromosome numbers.

**Extended Data Table 2.**
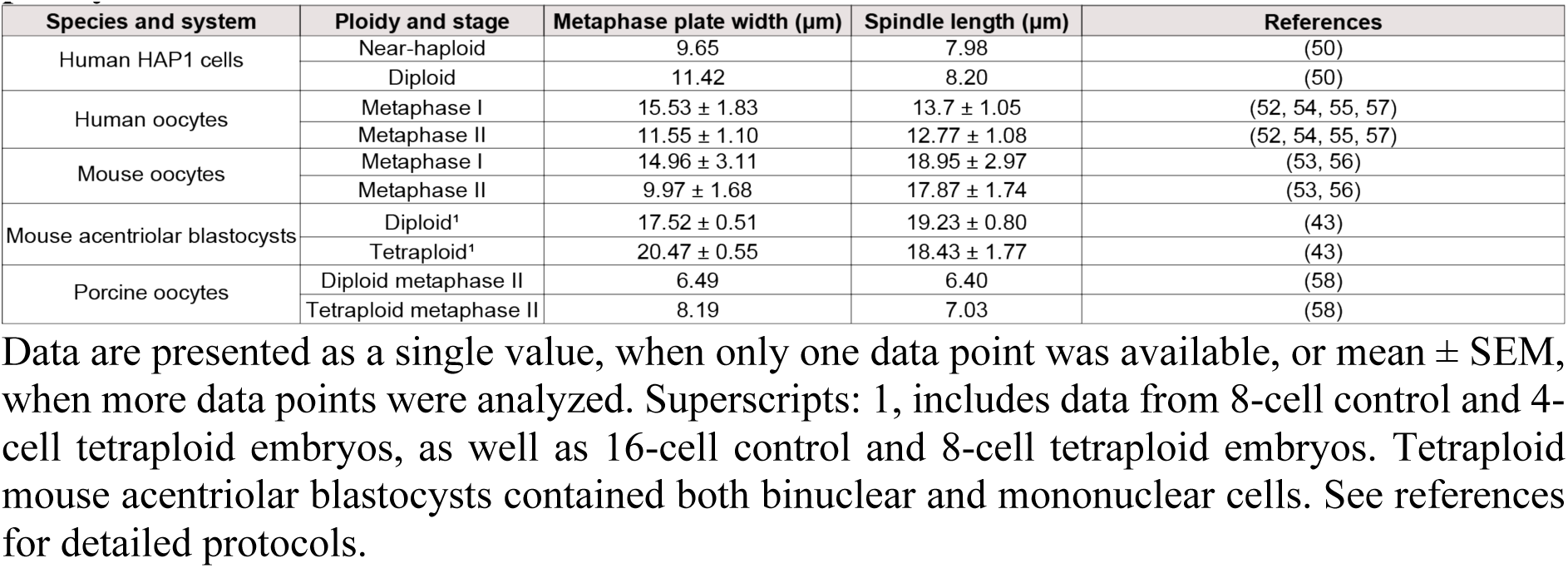
Measurements of metaphase plate width and spindle length from published images of acentriolar mitotic and meiotic spindles across different species and ploidy levels.

## Extended Data Movies 1-4

**Extended Data Movie 1. Diploid RPE1 cells and its derivatives of different ploidies during interphase and mitosis.** The combined movie shows live-cell imaging of the following: a diploid RPE1 cell (modal chromosome number = 46) expressing H2B-GFP; a hypotetraploid RPT1 cell (modal chromosome number = 80) expressing H2B-GFP; a newly generated mononuclear tetraploid cell with p53 knockdown (modal chromosome number = 92) expressing H2B-Dendra2; and a newly generated mononuclear hypooctaploid cell (modal chromosome number = 160) expressing H2B-GFP. Tetraploid and hypooctaploid cells were generated using the procedure detailed in Extended Data Fig. 2c and the Methods section. The movie was acquired by lattice light sheet microscopy with a time interval of 2 min. The movie plays at 7 frames per second. To ensure consistency, the first, second, and fourth segments include duplicate final frames of metaphase to match the number of frames in the third segment.

**Extended Data Movie 2. Acute compression of untreated spindles and spindles with an unreplicated genomes in U2OS cells.** The combined movie shows live-cell imaging of z stacks from hypertriploid U2OS cells expressing CENP-A-GFP (magenta), α-tubulin-mCherry (gray), and stained with Hoechst-DNA (cyan) under the following conditions: untreated control before compression (top left), mitosis with an unreplicated genome (MUG) before compression (bottom left), untreated control during compression (top middle), MUG during compression (bottom middle), untreated control after compression (top right), and MUG after compression (bottom right). MUG cells were generated using the procedure detailed in the Methods section. All z-stacks include the same number of slices, with the 0 point defined at the center of the mitotic spindle. The movie plays at 7 frames per second.

**Extended Data Movie 3. Acute reactivation of CENP-E activity in RPE1 cells.** The movie shows live-cell imaging of a diploid RPE1 cell expressing CENP-A-GFP and centrin-GFP (color-coded by depth using the 16-Color LUT in ImageJ) and labeled DNA (cyan) with SPY650-DNA (20 nM) overnight before imaging, following washout of a CENP-E inhibitor that was preincubated for 1 h. Experimental procedures are detailed in the Methods section. The movie is a maximum-intensity projection of the entire z-stack imaged by lattice light-sheet microscopy with a time interval of 10 s. The movie plays at 7 frames per second.

**Extended Data Movie 4. Nocodazole and osmotic shocks in RPE1 cells.** The movie shows live-cell imaging of diploid RPE1 cells expressing CENP-A-GFP and centrin-GFP (color-coded by depth using the 16-Color LUT in ImageJ) and labeled DNA (cyan) with SPY650-DNA (20 nM) overnight before imaging, following treatment with DMSO (left) 1 µM nocodazole (middle left), hypertonic solution (middle right), and hypotonic solution (right). Experimental procedures are detailed in the Methods section. The movie is a maximum-intensity projection of the entire z-stack with a time interval of 30 s. The movie plays at 7 frames per second.

